# Temporal Extracellular Vesicle Protein Changes following Intraarticular Treatment with Integrin α10β1-selected Mesenchymal Stem Cells in Equine Osteoarthritis

**DOI:** 10.1101/2022.09.30.510176

**Authors:** Emily J Clarke, Emily Johnson, Eva Caamano Gutierrez, Camilla Andersen, Lise C Berg, Rosalind E Jenkins, Casper Lindegaard, Kristina Uvebrant, Evy Lundgren-Åkerlund, Agnieszka Turlo, Victoria James, Stine Jacobsen, Mandy J Peffers

## Abstract

Equine osteoarthritis is a heterogeneous, degenerative disease of the musculoskeletal system with multifactorial causation, characterised by a joint metabolic imbalance. Extracellular vesicles are nanoparticles involved in intracellular communication. Mesenchymal stem cell (MSC) therapy is a form of regenerative medicine that utilises their properties to repair damaged tissues. Despite its wide use in veterinary practice, the exact mechanism of action of MSCs is not fully understood. The aim of this study was to determine the synovial fluid extracellular vesicle protein cargo following integrin α10β1-selected mesenchymal stem cell treatment in an experimental model of equine osteoarthritis with longitudinal sampling. Adipose tissue derived, integrin α10-MSCs were injected into the osteoarthritis afflicted joint after 18 days post surgery. Sixty-nine synovial fluid samples were collected via aseptic arthrocentesis at day 0, 18, 21, 28, 35, and 70. Synovial fluid was hyaluronidase treated and extracellular vesicles isolated using differential ultracentrifugation. Extracellular vesicles were characterised using the Exoview human tetraspanin chip. Extracellular vesicle concentration, surface marker identification, fluorescent microscopy and tetraspanin colocalization analysis was undertaken, in conjunction with nanoparticle tracking analysis. For proteomics, extracellular vesicle pellets were suspended in urea lysis buffer. Samples were reduced, alkylated and digested on SP3 beads with trypsin/LysC. A data independent acquisition mode was utilised for nano liquid chromatography tandem mass spectrometry analysis on a Triple TOF 6600 mass spectrometer. A total of 442 proteins were identified across all samples, with 48 proteins differentially expressed (FDR≤ 0.05) between control and osteoarthritis treated with MSCs.. The most significant pathways following functional enrichment analysis of the differentially abundant protein dataset were serine endopeptidase activity (p=0.023), complement activation (classical pathway) (p=0.023), and collagen containing extracellular matrix (p=0.034). To date this is the first study to quantify the global extracellular vesicle proteome in synovial fluid following MSC treatment of osteoarthritis. Changes in the proteome of the synovial fluid-derived EVs following MSC injection suggest EVs may play a role in mediating the effect of cell therapy through altered joint homeostasis and an improved phenotype.

## 1 Introduction

Osteoarthritis (OA) is a common disease of the joint, and is the cause of up to 60% of all lameness cases in horses (1). It is a progressive degenerative musculoskeletal pathology of synovial joints and results from an imbalance of catabolic and anabolic processes affecting cartilage and bone remodeling (2). This is associated with pain, reduced mobility and impaired welfare. OA is a complex heterogeneous condition with multiple causative factors, including mechanical, genetic, metabolic and inflammatory pathway activation, with a non-functional joint as the shared endpoint (3). In the disease there is a loss of articular cartilage, reduced elastoviscosity of synovial fluid, thickening of subchondral bone, joint space narrowing and osteophyte formation (4). Currently, OA is predominantly diagnosed based on clinical signs and radiographic imaging, capturing changes that are typical of the later disease stages. Treatment is symptomatic, with no current cure available to rescue the joint environment.

Biological cell-based therapies include autologous conditioned serum, platelet-rich plasma, and expanded or non-expanded mesenchymal stem cells (MSCs). These regenerative therapies have the potential to enhance repair of damaged tissues or organs (5). Mesenchymal stem cell (MSC) therapy most commonly uses cells derived from bone marrow or adipose tissue (6). They are highly proliferative, plastic-adherent, fibroblast-like cells that express CD44, CD90, CD105 and not express MHC class II or CD45 (7). MSCs are able to modulate and down-regulate immune system activity, reducing inflammatory cytokines associated with acute inflammation (8). MSC secreted factors are believed to have a critical role in the therapeutic efficacy. Cell-based therapeutics harbour inherent risks, such as cellular migration and uncontrollable division *in vivo*. Thus, there is a drive towards cell-free therapeutics that reflect the biophysical characteristic of ‘parent’ cells, whilst being less immunogenic (9).

Extracellular vesicles (EVs) are nanoparticles enveloped in a phospholipid bilayer membrane and are secreted by most mammalian cells. EVs facilitate intercellular communication through the paracrine of protein, lipid and nucleic acid cargo. EV subtypes, namely microvesicles and exosomes have been demonstrated to act in a protective manner, but also pathologically, dependent on the ‘parent’ cell phenotype and subsequent *in vivo* environment (10,11). Previous evidence has shown that EVs may at least partially-drive the therapeutic effect of MSC treatment. Zhang *et al*. (12) quantified the effect of bone marrow-derived MSC EVs in an osteochondral rat defect model. Exosomes increased cellular proliferation and infiltration in exosome-mediated cartilage repair and resulted in a regenerative immune phenotype, characterised by higher infiltration of CD163+ macrophages and a reduction in proinflammatory cytokines interleukin 1β (II.-1β) and tumour necrosis factor α (TNF-α)) (12). Tofiño-Vian *et al*. (13) found that in OA chondrocytes stimulated with IL-1β, EVs reduced the production of TNF-α, interleukin 6 (IL-6), and prostaglandin E2 (PGE2). Treatment of OA chondrocytes with EVs also led to a decrease in matrix metalloproteinase (MMP) activity and expression of MMP-13. Anti-inflammatory cytokines such as interleukin 10 (IL-10) and collagen type II were also upregulated (13).

In an equine model of OA, Hotham *et al*. (14) exposed chondrocytes stimulated with TNFα and IL1β to EVs obtained from autologous bone marrow-derived MSC-derived MSC. It was found that vesicles were taken up by the chondrocytes and had an anti-inflammatory effect on gene expression. A further study subjected equine chondrocytes to proinflammatory cytokines and then treated them *in vitro* with equine allogenic MSC-derived EVs (15). The MSC-EVs *in vitro* had an anti-inflammatory and anti-catabolic effect, with decreased expression of MMP-13. These observations are conserved across different species, such as in mice. Cosenza *et al*. exposed murine bone marrow-derived MSC exosomes and microvesicles to OA like murine chondrocytes. It was found that MSC-EVs could reinduce the expression of chondrocyte markers (type II collagen, aggrecan) while inhibiting catabolic (MMP-13) and inflammatory (nitric oxide synthase; iNOS) markers in murine chondrocytes. Additionally, it was found that exosomes and microvesicles could act in a chondroprotective manner by inhibiting apoptosis and macrophage activation (16).

It is paramount to determine the mechanism of therapeutic action of MSCs and determine the contribution of secreted factors such as EVs in rescuing the OA phenotype, with the aim of producing a more targeted treatment. We demonstrate that that intraarticular injection of MSCs affects protein cargo of synovial fluid EVs in an equine model of OA, decreasing the expression of proteins associated with pathways relating to OA pathogenesis.

## 2 Materials and Methods

### 2.1 Equine Carpal Osteochondral Fragment Model Induction

The study was approved by the Danish Animal Experiments Inspectorate (#2020-15-0201-00602) as well as the Ethical Committee of the University of Copenhagen (project no 2020-016). OA was induced using a carpal osteochondral fragment-exercise model in a total of 8 standard bred trotters6 trotter (mares, 4 to 7 years of age). The model was previously described by McIlwraith *et al*..(1) at Colorado State University. OA was induced in the left carpus of all horses and the right carpus was sham operated on to serve as control. Horses were premedicated with a combination of romifidine 6 mg/100 kg (Sedivet^®^Vet3, Boehringer Ingelheim Vetmedica, Missouri, United States), acepromazine 3 mg/100kg (Plegicil Vet4, Boehringer Ingelheim Vetmedica, Missouri, United States), atropine sulphate 0.5 mg/100kg (Atropin5, Aguettant Ltd, Bristol, United Kingdom), and butorphanol 3 mg/100 kg (Dolorex^®^6, Ag Marin Pharmaceuticals, United States). Anesthesia was induced with ketamine 2.5 mg/kg (Ketador Vet7, Richter Pharma AG, Oberosterreich, Austria) and midazolam 4 mg/100 kg (Midazolam “Accord”8, Accord-UK Ltd, Barnstaple, United Kingdom). The horses were placed in dorsal recumbency and anaesthesia maintained with isoflourane (Vetflurane9, Virbac, Carros, France). Perioperatively the horses received flunixin meglumine 1.1 mg/kg (Finadyne10, MSD Animal Health, New Jersey, United States), penicillin 22.000 IU/kg (Benzylpenicillin PanPharma11, Brancaster Pharma, Surrey, United Kingdom), and gentamicin 6.6 mg/kg (Genta-Equine12, Dechra Veterinary Products, Shrewsbury, United Kingdom). Arthroscopic portals were made in the right carpus and the carpal joint was inspected for abnormalities. In the left carpus an osteochondral “chip” fragment was made with an 8mm curved osteotome in the dorsal margin of the third facet of the distal surface of the radial carpal bone at the level of the medial plica.

The fragment remained attached to the plica. The debris was not flushed from the joint. The horses were stall rested for 14 days after surgery. From day 2 following surgery horses were walked by hand every day. Treadmill exercise was initiated on day 14 after surgery. The horses were exercised 5 days a week for 8 weeks through the following program: 2 min slow trot 16-19 km/h (4.4-5.3m/s), 2 min fast trot 32 km/h (9m/s), 2 min slow trot 16-19 km/h (4.4-5.3m/s). From day 14 the horses were allowed free pasture exercise every day. At day 18 MSCs were injected intraarticularly into the left joint only with the osteochondral fragment. Horses were humanely euthanized at the end of the study and both joints underwent both gross and histological examination.

Aseptic arthrocentesis was conducted on osteochondral chip joints and sham control joints at specific time points: day 0, 18, 21, 28, 35 and 70, as shown in Table 1 The synovial fluid (SF) was centrifuged at 2540*g* at 4□C for 5 minutes and then aliquoted into Eppendorf tubes, and snap frozen. After completion of collection, samples were transported on dry ice to the University of Liverpool, and stored at −80□C.

**Table 1.**
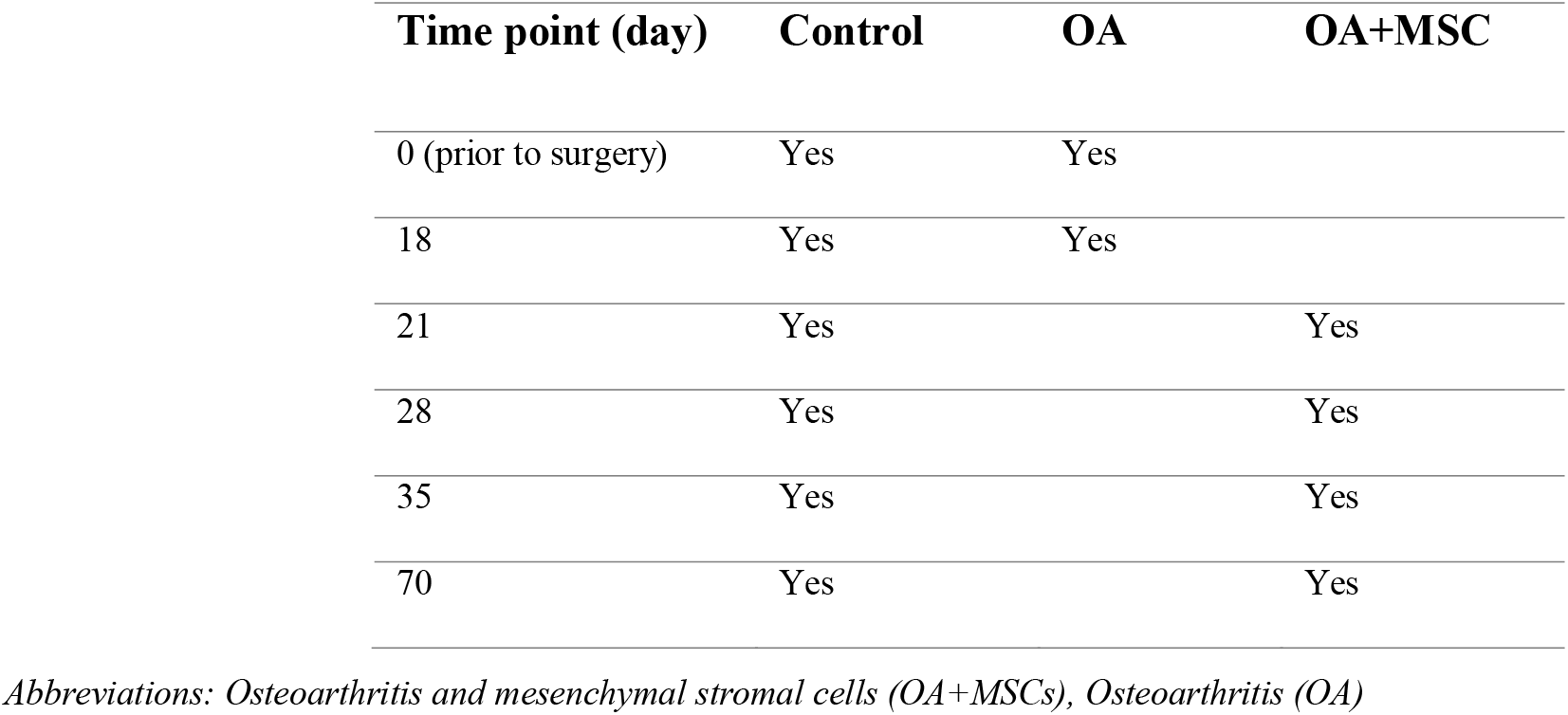
An overview of experimental groups and the longitudinal time points for SF collection.

### 2.2 Mesenchymal Stem Cell Therapy

Equine MSCs were isolated from adipose tissue from a 7-year-old healthy gelding. Briefly, the adipose tissue was digested with collagenase, adipose cells were removed, and the stromal vascular fraction was isolated and expanded in culture. Cells were cultured to passage 3, and MSCs were then selected for high expression of integrin α10β1 (integrin α10-MSCs) by magnetic-activated cell sorting using a specific biotinylated integrin α10 monoclonal antibody (Xintela, Sweden) and anti-biotin microbeads (Miltenyi, North Rhine Westphalia, Germany). Only MSCs with a high expression of integrin α10β1 (integrin α10-MSCs) were used. MSCs were subsequently washed in culture medium, reseeded and expanded for a further passage, then frozen in dimethyl sulfoxide cryopreservation medium (Cryostor, BioLife Solutions, Washington, United States), in liquid nitrogen until use.

On day 18 all six horses were treated with 20 ×10^6^ integrin α10-MSCs immediately after synovial sampling. The integrin α10-MSCs were thawed in a sterile water bath at 37°C, aspirated into a syringe through a 14G canula at a slow pace, and injected into the carpal joint of the OA-induced leg through a 20G canula over a minimum of 10 seconds. The horses were stall rested for 2 days following treatment.

### 2.3 Sample Preparation

One ml of SF per sample was spun in a benchtop centrifuge (Eppendorf non-refrigerated centrifuge 5420) at rpm1400rpm for 10 min. The supernatant was removed and treated with hyaluronidase (1 μg/ml) by incubation at 37°C for 1 hour. SF samples were then spun at 1000 x g for 5 minutes, and the supernatant was collected.

### 2.4 EV Isolation – Differential Ultracentrifugation

Equine SF samples (200 μl) underwent differential ultracentrifugation (dUC) in order to isolate EVs. Samples were subjected to a *300g* spin for 10 min, 2000*g* spin for 10 min, 10,000*g* spin for 30 min in a bench top centrifuge at room temperature. Samples were then transferred to Beckman Coulter thick wall polycarbonate 4 mL ultracentrifugation tubes, and centrifuged at 100,000*g* for 70 min at 4°C (Optima XPN-80 Ultracentrifuge, Beckman Coulter, California, USA) in a 45ti fixed angle rotor, with the use of a 13 mm diameter Delrin adaptor. Supernatant was removed and sample pellets were resuspended in 50 μl of filtered phosphate buffered saline (PBS) (Gibco™ PBS, pH 7.4 - Fisher Scientific, Massachusetts, USA), resulting in 50 μL SF-EV (SF-EV) per sample.

### 2.5 Nanoparticle Tracking Analysis

Nanoparticle tracking analysis (NTA) was used to quantify EV concentration and size in all samples, using a NanoSight NS300 (Malvern Panalytical, Malvern, UK). Samples were prepared as previously described (17).

### 2.6 EV characterisation

The ExoView platform (NanoView Biosciences, Malvern Hills Science Park, Malvern) was used to determine EV concentration, surface marker identification (CD9, CD81 and CD63) and to perform fluorescent microscopy and tetraspanin colocalisation analysis on selected samples. We had previously tested equine samples on both the human and murine chips and demonstrated the human chips were more compatible (data not shown). The ExoView analyzes EVs using visible light interference for size measurements and fluorescence for surface protein profiling. Samples were analysed as previously described (18).

### 2.7 EV Protein Extraction

EV pellets were suspended in 200 μl of urea lysis buffer (6M Urea (Sigma-Aldrich, Dorset, United Kingdom), 1M ammonium bicarbonate (Fluka Chemicals Ltd., Gillingham, UK) and 0.5% sodium deoxycholate (Sigma-Aldrich, Dorset, United Kingdom)). Samples were sonicated at 5 μm for 3x 10 seconds per sample, with 1-minute rest on ice between each sonication round.

### 2.8 SDS-PAGE and protein staining

Sodium dodecyl sulphate–polyacrylamide gel electrophoresis (SDS-PAGE) was used to separate proteins from EV protein extract. 7.5 ul of 2x Novex™ Tris-Glycine SDS Sample Buffer (ThermoFisher Scientific, Paisley, UK), supplemented with 8% of 2-Mercaptoethanol (Sigma-Aldrich, Dorset, UK), was added to 7.5ul of sample SF-EV protein lysate. Samples were mixed and heated at 100°C for 10min to denature proteins then placed on ice. A NuPAGE™ 4 to 12%, Bis-Tris gel (ThermoFisher Scientific, Paisley, UK) was placed in the electrophoretic tank and the tank was filled with 1x NuPAGE^®^ MES Running Buffer (ThermoFisher Scientific, Paisley, UK) (diluted from the 20x stock in ultrapure water). Samples were loaded onto the gel alongside the Novex™ Sharp Pre-stained Protein Standard ladder (ThermoFisher Scientific, Paisley, UK). Gels were run at 100V until completion of electrophoresis and visualised using colloidal coomassie blue (Thermofisher Scientific, Paisley, UK) according to manufacturer’s guidelines.

### 2.9 In-solution Digestion

95 μL of lysed and sonicated equine SF-EV were treated with 5mM dithiothreitol (DTT) (Sigma-Aldrich, Dorset, UK) 100 mM at 60°C and 1000rpm for 30 minutes. Iodoacetamide (Sigma-Aldrich, Dorset, UK) was then added to a final concentration of 20mM and the samples were incubated at room temperature in the dark for 30 minutes. Samples were quenched with 5mM DTT, and incubated at room temperature for 15 minutes. 12μl hydrophilic and hydrophobic magnetic carboxylate SpeedBeads (SP3 beads, total of 12μL) (Cytiva, Massachusetts, United States) were added to each samples, followed by 120 μL ethanol (Sigma-Aldrich, Dorset, UK). Samples were then incubated at 24°C and 1000rpm for 1 hour. The beads were separated from samples using a magnetic stand and were washed three times with 180μL 80% ethanol. They were resuspended in 100 mM ammonium bicarbonate (Fluka Chemicals Ltd., Gillingham, UK4 μg). Trypsin/LysC (2.4μg) (Promega) was added to each sample. Samples were placed in a sonicator bath and sonicated for 30 seconds to disaggregate the beads before being incubated overnight at 37°C and 1000 rpm.1000rpm. Beads were removed from the samples using the magnetic stand and the supernatants were acidified by the addition of 1 μl trifluoroacetic acid, (Sigma-Aldrich, Dorset, UK). Samples were then desalted using an Agilent mRP-C18 column, dried in a SpeedVac and resuspended in 0.1% formic acid. The UV absorbance measured during desalting was used to normalize the loading for mass spectrometry analysis with a final volume of 5μL being loaded on the nano-LC column.

### 2.10 Data-dependent acquisition (DDA) for generation of an equine SF EV spectral library

Equine SF was pooled from the metacarpophalangeal and carpal joint of control joints as well as OA, and MSC treated joints resulting in a total of 11 ml SF. This was hyaluronidase treated (1μg/ml) for 1 hour at 37°C, as outlined in section 2.2. EVs were isolated using dUC, as outlined in section 2.4. The EV pellet was then reconstituted in 200 μl of urea lysis buffer. The samples were digested with trypsin/LysC for 3 hours at 37°C, the concentration of urea was reduced to 1M, and incubation was continued overnight at 37°C. Samples were fractionated on a PolySULFOETHYL a strong cation exchange column and 20 fractions were desalted, dried and resuspended in 0.1% formic acid. Aliquots were loaded onto an Eksigent nanoLC 415 (Sciex, Macclesfield, United Kingdom) equipped with a nanoAcquity UPLC Symmetry C18 trap column (Waters, Massachusetts, United States of America) and a bioZEN 2.6μm Peptide XB-C18 (FS) nanocolumn (250mm x 75μm, Phenomenex, Macclesfield, United Kingdom). A gradient from 2-50% acetonitrile /0.1% formic acid (v/v) over 120 min at a flow rate of 300 nL/min was applied. Data-dependent acquisition was performed using nano liquid chromatography tandem mass spectrometry on a Triple TOF 6600 (Sciex, Macclesfield, United Kingdom) in positive ion mode using 25 MS/MS per cycle (2.8s cycle time), and the data were searched using ProteinPilot 5.0 (Sciex, Macclesfield, United Kingdom) and the Paragon algorithm (SCIEX) against the horse proteome (UniProt *Equus cabullus* reference proteome, 9796, May 2021, 20,865 entries). Carbamidomethyl was set as a fixed modification of cysteine residues. The data were also searched against a reversed decoy database and proteins lying within a 1% or 5% global false discovery rate (FDR) were included in the library. Proteins were analysed using FunRich.

### 2.11 Data-independent acquisition proteomics

A data-independent proteomic approach was utilised in the form of Sequential Windowed Acquisition of all theoretical fragments (SWATH) (19). Aliquots of 5μL containing equal quantities of peptides were made up to a volume of 5 μL with 0.1% formic acid and data were acquired using the same 2h gradient as the library fractions. SWATH acquisitions were performed using 100 windows of variable effective isolation width to cover a precursor m/z range of 400-1500 and a product ion m/z range of 100-1650. Scan times were 50ms for TOF-MS and 36ms for each SWATH window, gave a total cycle time of 3.7 seconds. Retention time alignment and peptide/protein quantification were performed by Data-Independent Acquisition by Neural Networks (DIA-NN) (20), using the same reference horse proteome as above to reannotate the library. A precursor FDR of 1%, with match between runs and unrelated runs was selected. The mass spectrometry proteomics data were deposited to the ProteomeXchange Consortium via PRIDE (21) (reference PXD035303).

### 2.12 Statistical Analysis

Nanoparticle tracking analysis data was analysed using non-parametric tests. Kruskal Wallis test was performed for concentration and size with FDR correction Benjamini-Hochberg (B-H), followed by a Mann Whitney test per time point. Exoview data was analysed using T-tests following parametric evaluation in GraphPad Prism 9.0 All statistical analysis of proteomic data was carried out using the R statistical programming environment (22), unless stated otherwise. Proteins with complete values were log2 transformed and used for downstream analysis. Data was QC and batch effect detected and corrected via ComBat prior to Principal Component Analysis (PCA) and further visualisations were log2 transformed. (23). The package lme4 (22) was used to fit linear (23). Linear mixed models (LMMs) were fit to the log-transformed data for each protein to determine the effect of MSC treatment on OA horse joints compared to the control joints over time to determine differences related to treatment, time and their interaction. Significant results were adjusted for FDR by the B-H method and considered significant at 5% FDR. Pairwise comparisons between the treatment and control were carried out at each time point using the emmeans package with the Kenward-Roger method (24). All graphical representations were undertaken using the package ggplot2 (25). Functional classification and enrichment analyses were performed using the clusterProfiler package (26). The proteins were annotated with GeneOntology (GO) terms using the UniProtKB ID Mapping tool. UniProt keywords. UniProtOver-representation analysis (ORA) was carried out for GO terms UniProt using the enricher function from the clusterProfiler package. The foreground was all the proteins that passed FDR, the background was all the processed proteins after missing values had been removed, i.e., all the proteins that were subjected to statistical analysis. Each term was required to have a minimum of three observed proteins annotated to it and an adjusted p-value <0.05.

## 3 Results

### 3.1 Equine Carpal Osteochondral Fragment Model

The carpal osteochondral fragment-exercise model used in this experiment has been shown to result in an OA phenotype with respect to clinical parameters, and responded to integrin α10-MSC treatment by reduced lameness and joint degradation, as shown by Andersen *et al*. 2022 (submitted to Stem Cell Research and Therapy).

### 3.2 Nanoparticle Tracking Analysis (NTA) to quantify EV size and concentration

Total particle concentration and size characterisation was performed using NTA. NTA determined the average SF-EV sample concentration between control, OA and OA+MSCs across specific time points, specifically quantifying all nanoparticles within the sample (Figure 1A). No significant difference was observed in EV concentration between experimental groups, however a significant difference in EV size was found irrespective of time when comparing control and OA (p=0.02) and control compared to OA+MSCs (p=0.02). Specifically, EV size was significantly different between control and OA at day 18 before MSC treatment (p=0.02) (Figure 1B). Results were suggestive of a heterogeneous population of nanoparticles.

**Figure 1.**
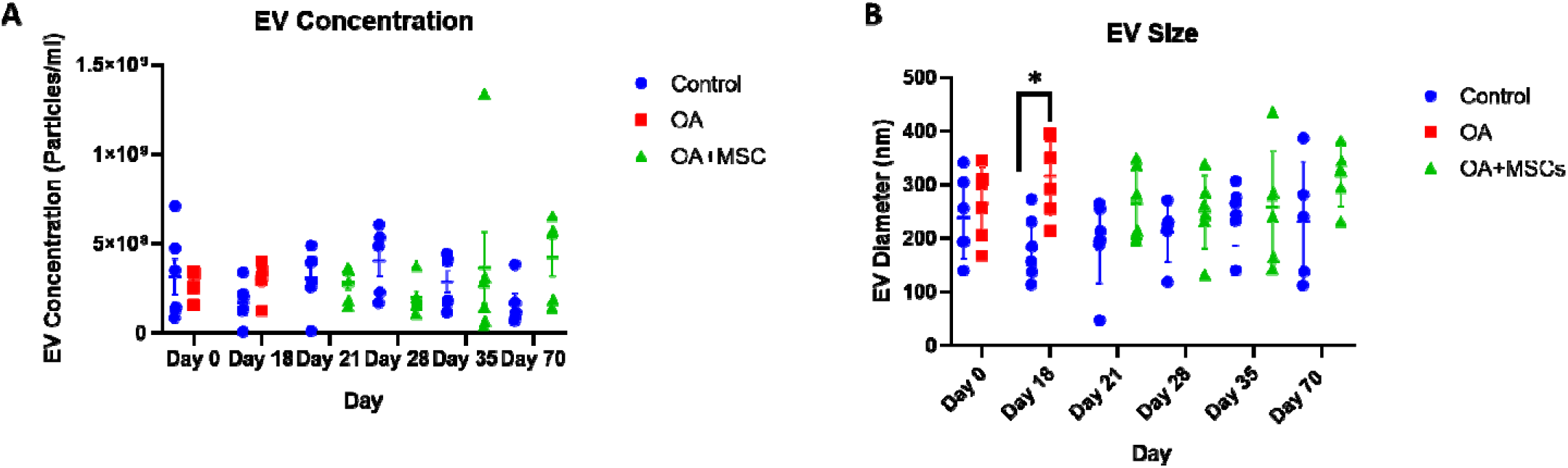
Size and concentration of synovial fluid-derived nanoparticles. Nanoparticle tracking analysis was undertaken using a Nanosight NS3000. All error bars are standard error of the mean (SEM). Statistical analysis undertaken in GraphPad Prism 9.0 using Kruskal Wallis Tests with FDR correction and Mann Whitney Tests within time points. (P<0.05, *; P<0.01 **: p<0.001, ***, p<0.0001, ****). (1A) EV concentration and (1B) EV size.

### 3.3 Exoview assay characterises equine synovial fluid extracellular vesicles, including morphology and surface tetraspanins

In addition to NTA, representative EV samples were characterised using the human exoview tetraspanin chip assay. This assay specializes in characterizing the exosome subpopulations of EVs. Control at day 0, OA and control at day 18 and OA+MSCs and control at day 70 were compared. OA and OA++MSCs groups had a significantly higher concentration of EVs when compared with controls. For CD9 expression, control had 4.63 ×10^3^ particles, OA had 21.91×10^3^ particles, and OA with MSCs had 15.97 ×10^3^ particles. Similarly, for CD81, control had 3.41 ×10^3^ particles, OA had 17.23 ×10^3^ particles and OA with MSCs had 12.48 ×10^3^ particles (Figure 2). CD63 was not reported due to low particle counts for this tetraspanin; this has been attributed to poor protein homology between equine and human CD63 tetraspanins. EVs were visualised between groups with fluorescent microscopy, highlighting tetraspanin expression and EV morphology (Figure 3).

**Figure 2.**
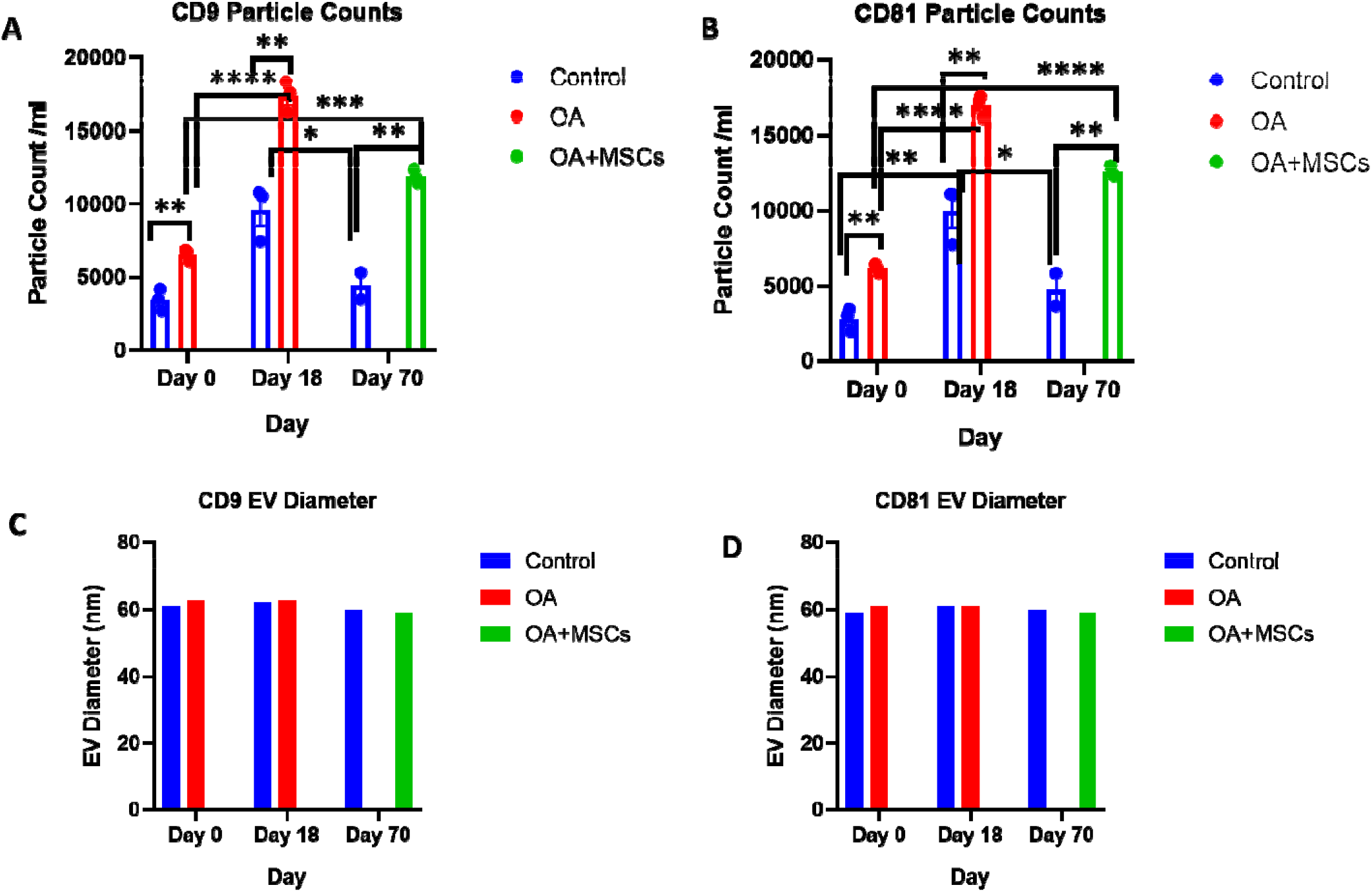
Sizing and enumeration of synovial fluid-derived EVs. All data was adjusted for dilution of the sample onto the chip. Shown is the average representing mean of three technical replicates th at were run for each sample. Particle numbers were quantified by the number of particles in a defined area on the antibody capture spot. All bars are mean and standard error mean. A. CD9, and B. CD81-positive particles following probing with fluorescent tetraspanin antibodies. C. Sizing of CD9 and D. CD81 labelled EVs, normalised to MIgG control. Limit of detection was 50-200 nm. Statistical analysis undertaken in GraphPad Prism 9.0 using T-tests following parametric evaluation (P<0.05,*; P<0.01 **: P<0.001, ***, P<0.0001, ****).

**Figure 3.**
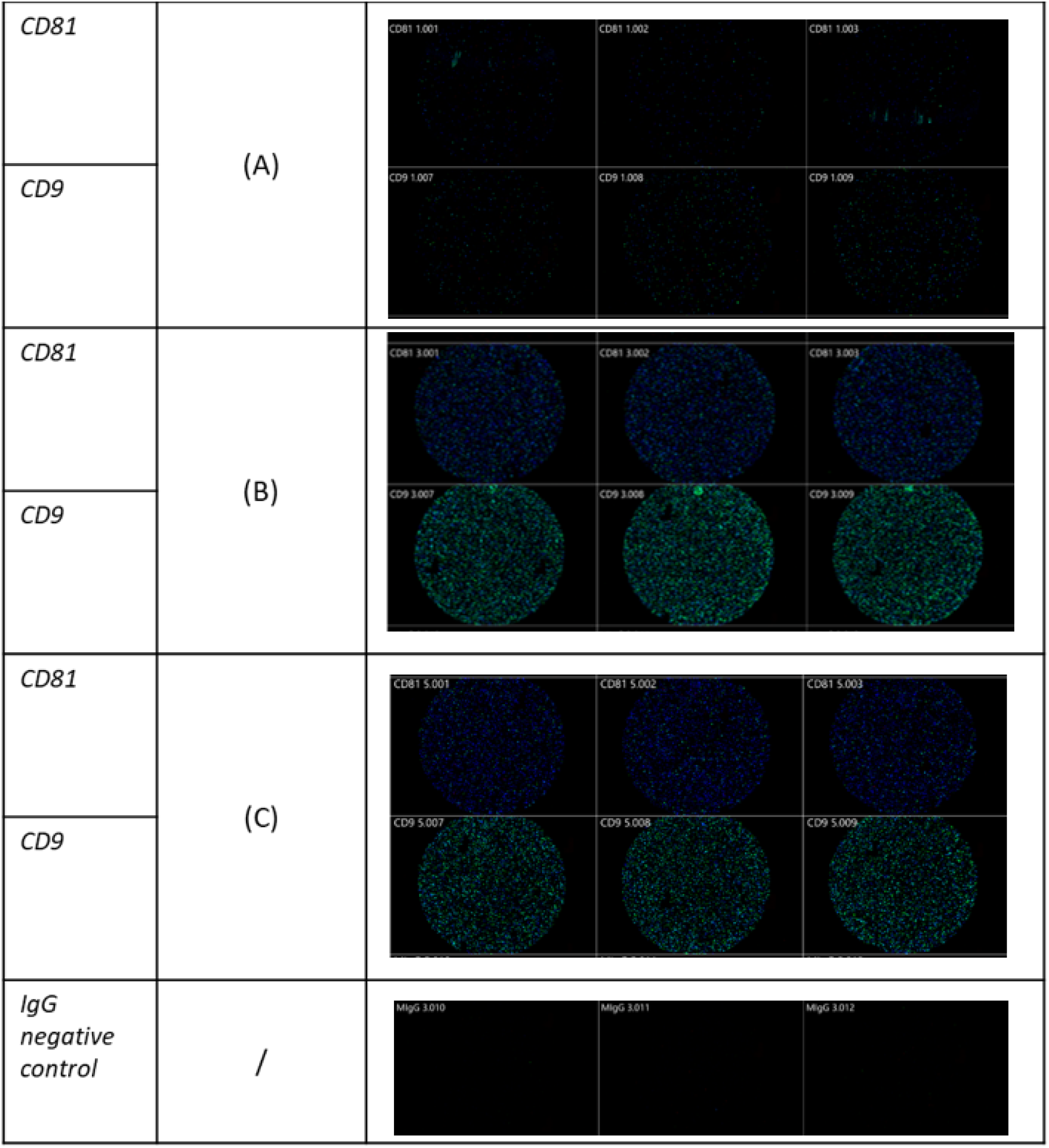
Visualization of SF-derived EVs from control, osteoarthritic (OA) and OA +MSCs joints using Exoview at selected time points. A fluorescent image of a representative spot is shown for each sample comparing (A) control, (B) OA and (C) OA +MSCs with colour denoting surface tetraspanin positive identification (blue-CD9, and green -CD81).

### 3.4 Spectral library for equine synovial fluid extracellular vesicles

Mass spectrometric analysis of the SF-EV pooled sample identified 2271 proteins, mapping at least one unique peptide. Of these proteins 2047 were identified and mapped to GO Cellular Component terms using FunRich. Proteins were attributed to various cellular components, including extracellular space and exosomes, both p ≤ 0.001, as shown in Figure 4. This library was then used to identify the proteins present within the individual samples.

**Figure 4.**
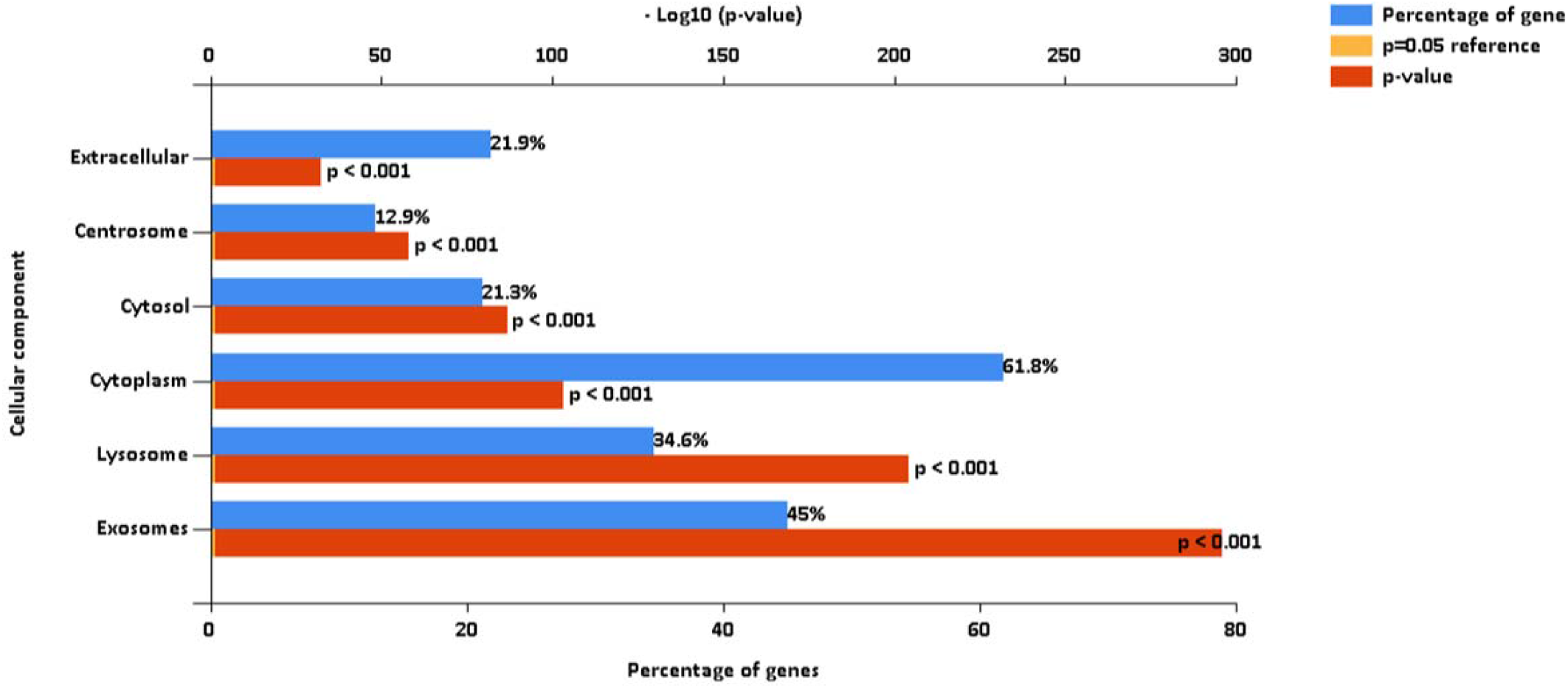
FunRich analysis output, after mapping 2047 proteins to GO Cellular Component terms. These proteins were identified from a pooled sample of equine synovial fluid (11ml) used to generate the SF-EV spectral library for this study. SF was sourced from healthy, OA and OA+MSC treated joints in order to encapsulate all potential proteins that may be present across all experimental groups.

### 3.5 A multivariate approach identified a time-dependent difference between disease stages pre and post-treatment

A total of 442 proteins were identified across all samples submitted for SWATH -MS analysis. Multi-level PCA (mPCA) was carried out using the MixOmics R package principal component analysis (mPCA). PCA normally assumes the variables are not correlated. However, this study employed a repeat-measures design on the same horses. To account for the intraclass correlation between horses mPCA was employed. The first two principal components accounting for 54% of the variance were associated with the biological effect and demonstrated that the control and OA+MSC samples clustered by treatment (Figure 5). The later OA+MSC time points (day 70) appeared to cluster together with the control samples, reflecting a return to protein expression levels comparable to the healthy controls.

**Figure 5.**
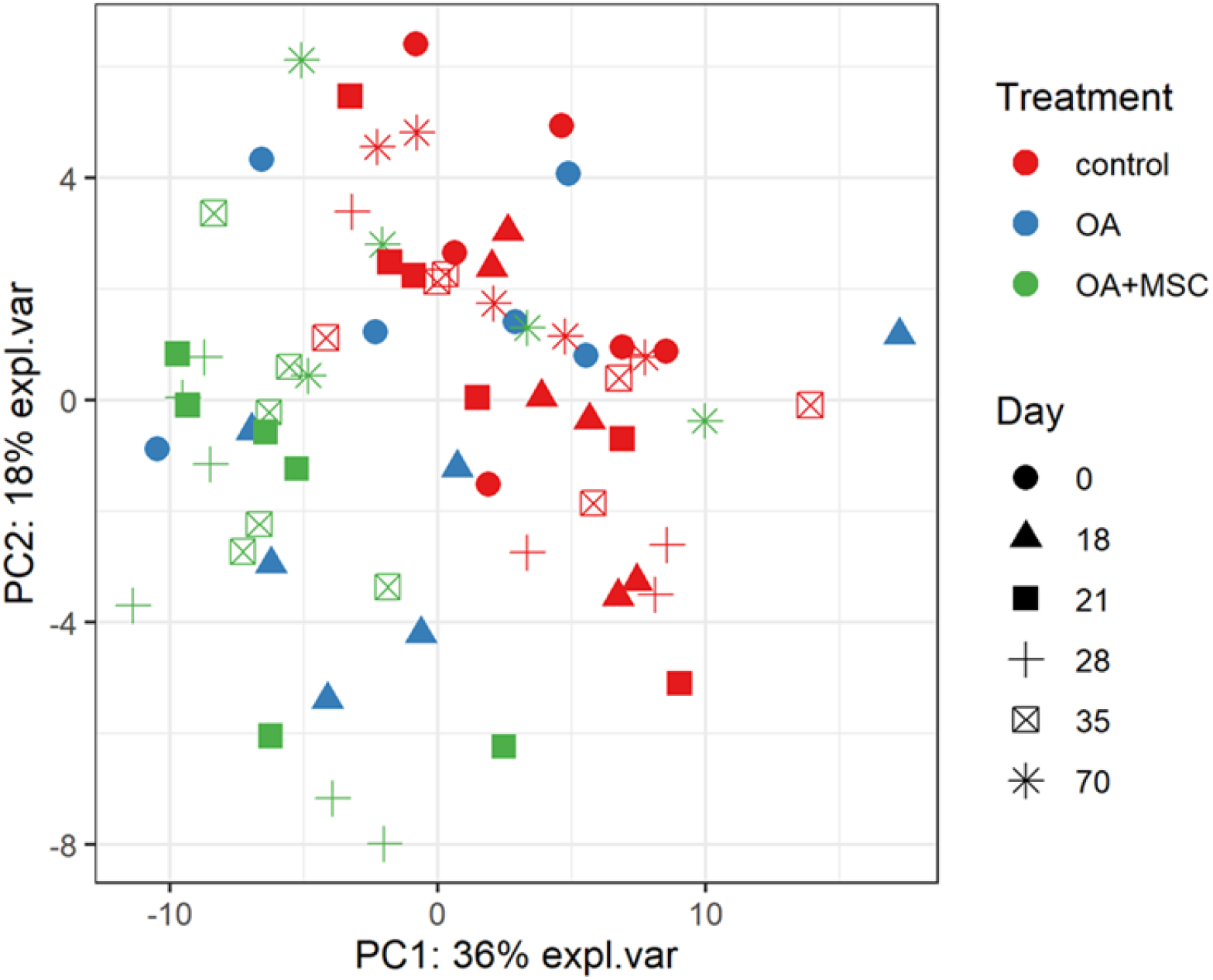
Multi-level PCA (mPCA). The first two principal components are plotted, accounting for ~54% of variance. samples based on SWATH-MS. Each plotted point represents a horse, which are coloured by their treatment and shaped by the day of the study.

### 3.6 Longitudinal differential expression highlights 6 key pathways related to disease progression and treatment

Differentially expressed (DE) proteins were identified with the application of a linear mixed model with regressed factors comparing protein expression between experimental groups across time. Statistical significance was attributed to any protein with an FDR corrected p value (B-H) of p≤0.05 meeting a minimal 95% confidence interval. Of the 442 proteins identified, 48 were presentat significantly different levels (p≤ 0.05) between control and OA+MSCs regardless of time. Interestingly, there were no proteins DE in SF-EVs between control and OA at any time point. The 10 proteins with the most statistically significant altered levels are shown in Table 2, and all 48 can be found in Supplementary Table 1. Figure 6 shows proteins attributed to the serine endopeptidase (p=0.02) and complement pathways (p=0.03), included hyaluronan binding protein 2 (p=0.04) (Figure 6A), complement subcomponent C1r (p=0.03) (Figure 6B), CD5 (p=0.01) (Figure 6C), complement factor D (p=0.03) (Figure 6D), C2 (p=0.04) (Figure 6E), and C1 (p=0.04) (Figure 6F). Other proteins attributed to the serine endopeptidase pathway include haptoglobin (p=0.03), HtrA1 serine peptidase (p=0.03) and complement factor B (p=0.04). Proteins mapped to the collagen containing extracellular matrix (p=0.02) included cartilage oligomeric matrix protein (p=0.001), microfibril associated protein 4 (p=0.001), thrombospondin 4 (p=0.003), retinoic acid receptor responder protein 2 (p=0.03), periostin (p=0.03), EGF containing fibulin extracellular matrix protein 1 (p=0.03), and cartilage intermediate layer protein (p=0.04). (Supplementary Figure 1). The third most significant pathway was identified as complement activation classical pathway, with the following proteins attributed to it; two uncharacterized proteins, C9 (p=0.01), C8A (p=0.02), C1r (p=0.03), C7 (p=0.03), C2 (p=0.04), and C1 (p=0.04) (Supplementary Figure 1). Across all figures (Figures 6A–4F) protein expression was significantly different at day 21, 28 and 35 when compared to control. All protein expression returned to baseline control by day 70.

**Table 2.**
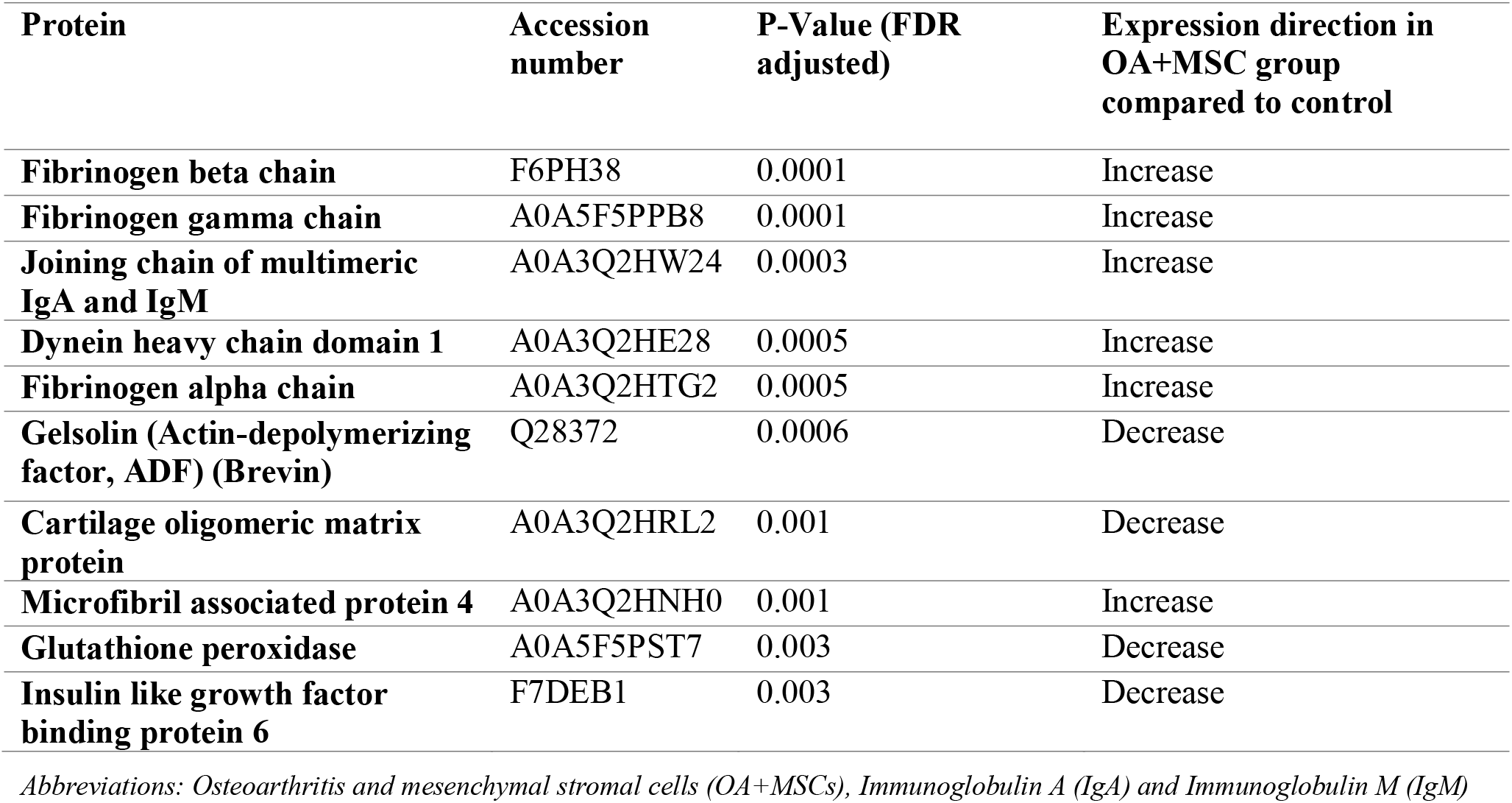
The top 10 differentially expressed (P<0.05) proteins accounting for treatment and timepoint. Treatment and time were included as main effects, along with a treatment-by-time interaction term.

**Figure 6.**
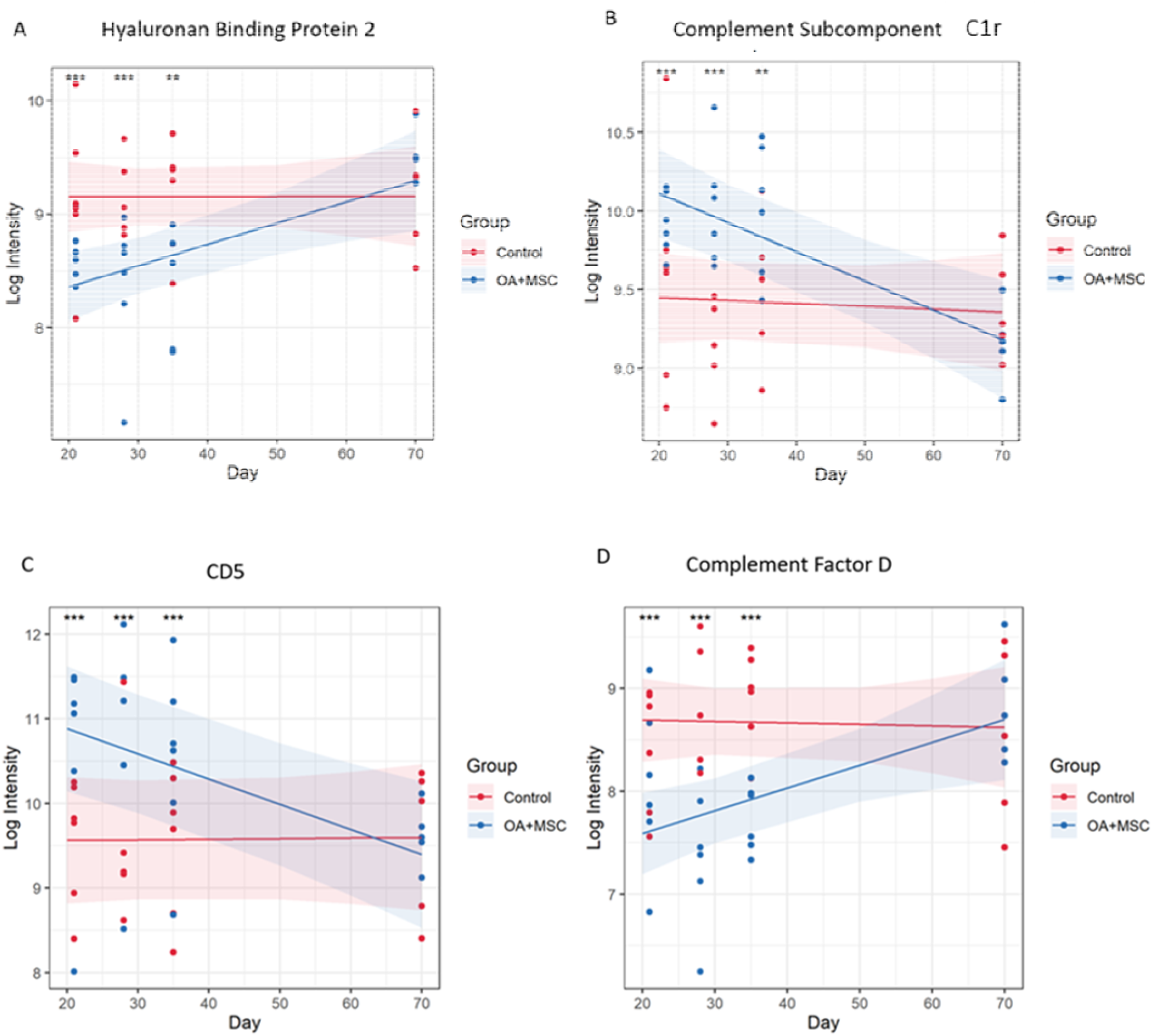
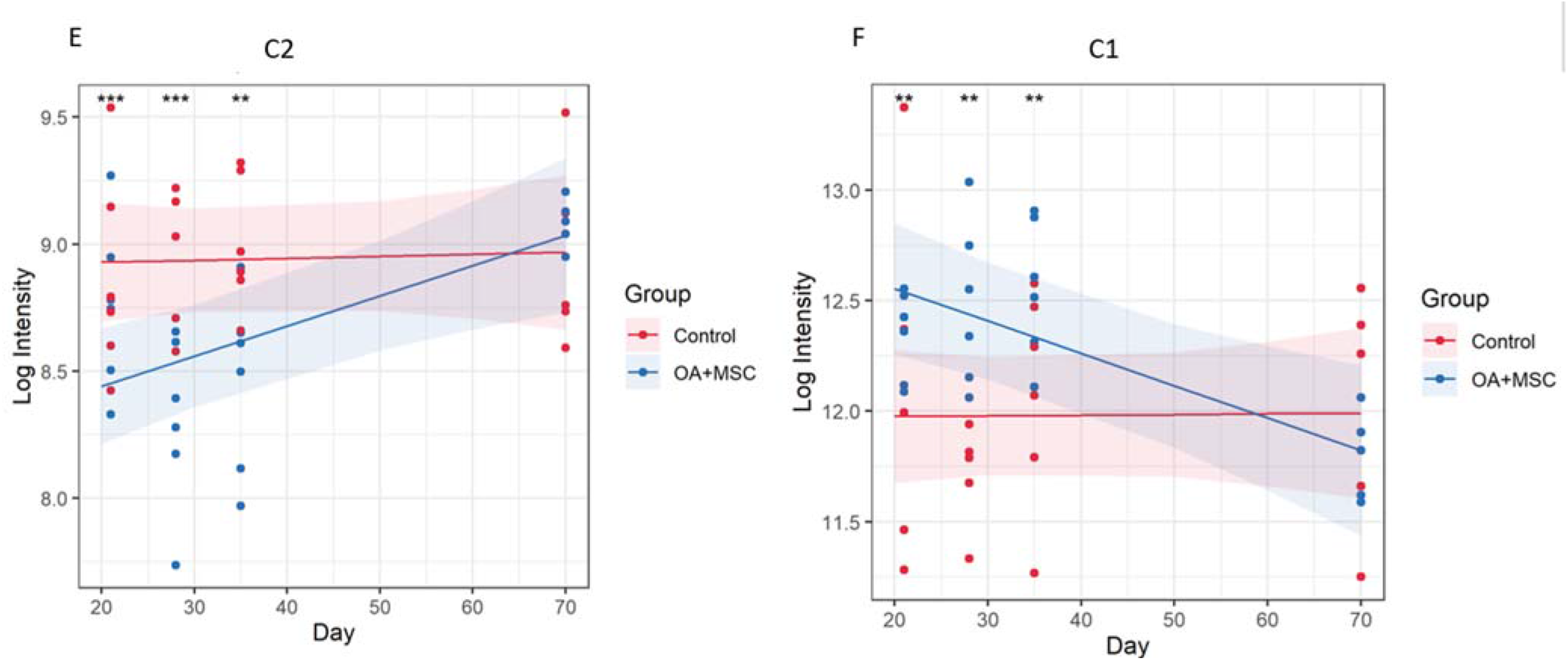
Protein expression changes in response to MSC treatment in OA joints vs control. (A) Hyaluronan binding protein 2, (B) Complement subcomponent C1r, (C) CD5, (D) Complement factor D, (E) C2, (F) C1. The models were fitted using the lmerTest implementation of lme4.48 proteins had a significant group, time, or group:time effect after FDR correction. For each protein with a significant effect the model was plotted using the effects and ggplot2 packages. The fitted model is shown as a line for the OA+MSC (blue) and control (red). The 95% confidence intervals for each group are shown as a shaded area. The raw data is included as points. At each time point pairwise comparisons were carried out between the treatment and control using the emmeans package. Significance thresholds were as follows: (P<0.05, *; P<0.01 **: P<0.001, ***, P<0.0001, ****)

The lists of significant proteins were used to calculate functional enrichment with the aim of identifying relevant biological processes representative of the signature. Using an overenrichment analysis (ORA) approach on gene ontology (GO) annotations yielded six enriched pathways: serine endopeptidase activity (p=0.023), complement activation (classical pathway) (p=0.023), collagen containing extracellular matrix (p=0.034), protein polymerization (p=0.039), platelet aggregation (p=0.039), elastic fibre (p=0.039) (Figure 7).

**Figure 7.**
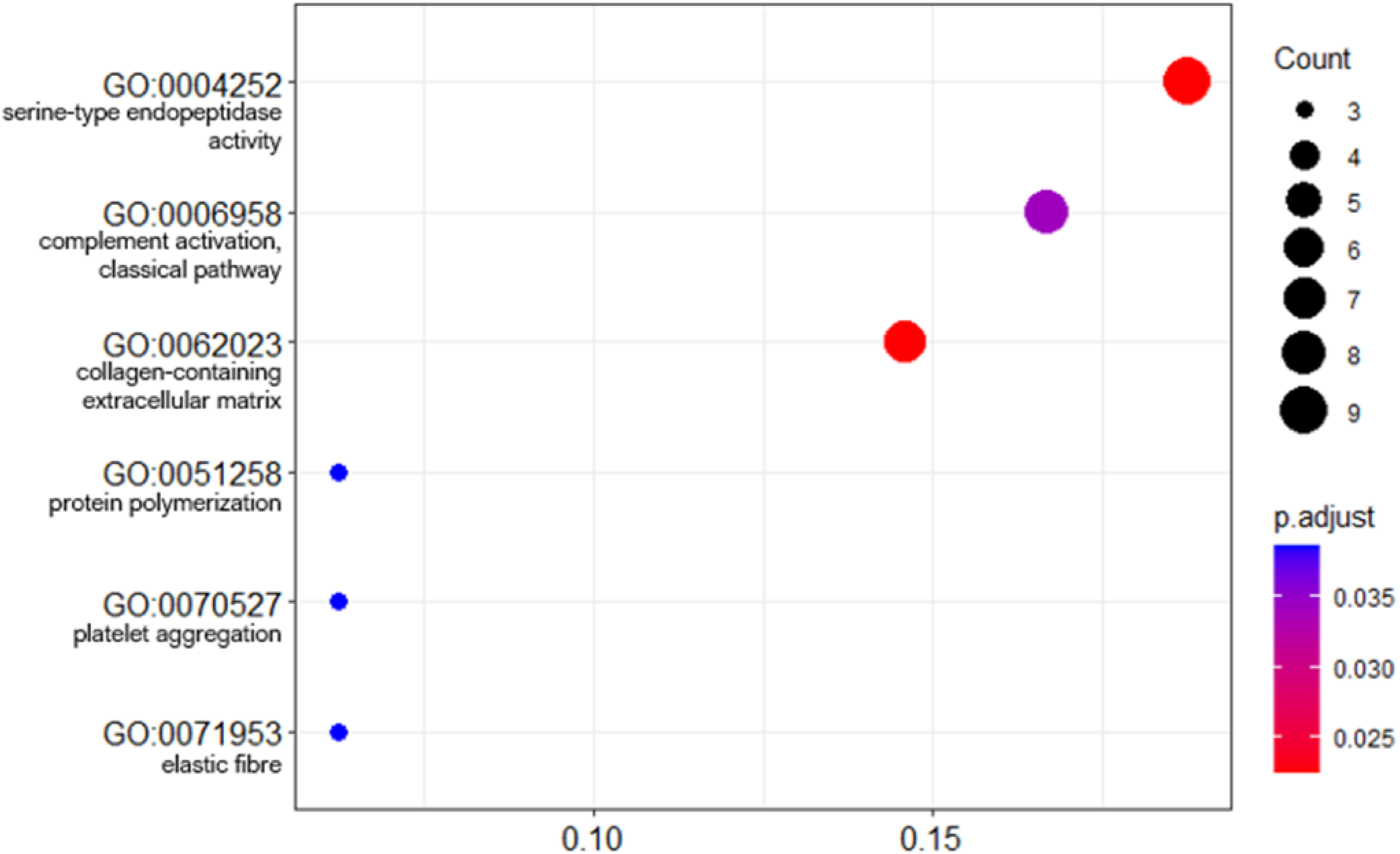
Dot plot of GO term enrichment analysis of differentially abundant proteins. The size of the dots indicates the number of proteins that mapped to that term. The x-axis is the protein ratio (number of proteins that map to the term divided by the total number of significant proteins). The dots are shaded by adjusted p-values (BH method).

## 4 Discussion

This study aimed to determine the EV protein cargo following integrin α10β1-selected mesenchymal stem cell (integrin α10-MSC) treatment in an experimental model of equine OA. We identified a global change in the EV proteome, and identified a possible mechanism of MSC therapy. A time-dependent change in the EV protein cargo was also observed, suggestive of a time associated therapeutic effect. To our knowledge this is the first study of its kind to quantify the global EV proteome *in vivo* after MSC treatment.

We used dUC to isolate EVs following hyaluronidase treatment of SF. Hyalaronidase was used in order to breakdown hyaluronic acid as its presence in SF increases viscosity making the biofluid difficult to handle. This pretreatment is known to increase EV yield (35). In addition, it has also been suggested that SF-derived EVs should be sedimented at a speed of at least 100,000*g* for optimal EV recovery, hence the decision to use this step within our isolation protocol (35). There are many EV isolation protocols with no standardized protocol agreed upon. Each isolation method results in different sample concentrations, purity, and EV profiles/ heterogeneity.

EVs were isolated from SF obtained from control and OA joints that were treated with MSCs. The EVs isolated were a heterogenous population derived from cells found in the intraarticular environment, such as synoviocytes and chondrocytes, as well as from the MSCs injected into the joint. Nanoparticle tracking analysis conducted on all samples across all time points found no changes in EV concentration with time. It should be noted that NTA quantifies all nanoparticles within a sample and includes lipoproteins, proteins, viruses, nanovectors and drug delivery systems (27). This accounts for the difference in EV concentration between our NTA analysis and exoview analysis. We had limited resources to enable Exoview of all samples, and so used the platform for a subset of samples. Exoview analysis specifically focusses on the exosomal population of EVs by using antibodies for surface tetraspanins such as CD9, CD81 and CD63. Exoview analysis demonstrated an increase in the number of exosomes after OA induction prior to MSC injection, converse to human studies, such as Mustonen *et al*. (2022) which identified no change in EV concentration between SF-EVs from healthy and human OA patients (28). Our results tentatively suggest that OA induction actively increases the number of exosomes. This could be due to the access of subchondral bone to the SF environment following production of an osteochondral fragment or from tissues within the joint as a response the formation of the fragment or both. However, our experimental design does not enable us to decipher this. A greater number of CD9+and CD81+ EVs were identified across all experimental groups using the exoview assay. CD9+ exosomes have been postulated to be a target for inflammatory regulation in specific pathologies (29). In addition, the increase count observed in OA+MSC groups compared with control could be attributed to its presence on hematopoietic cells, and the role of CD9 in regulating hematopoietic stem cells differentiation (29). Across all experimental groups the sham control remained significantly lower than the experimental group at all time points, suggestive of a minimal effect from the surgical opening of the joint in the sham control groups. In addition, highlighting that there was a limited systemic effect from the surgical induction of OA.

In this study, SF samples following OA induction and then following the addition of MSCs to the joint were available up to day 70 of treatment post-induction. Unfortunately, SF samples from OA joints without MSC treatment were not available for this study. This makes it difficult to be definitive about the source of the EVs in the SF following addition of MSCs. We believe these will be a combination of tissue-derived and MSC-derived EVs, resulting from MSC-to-cell interactions, cell-to-MSC interactions, and cell-to-cell interactions.

Post intraarticular injection of MSCs an increase in EV COMP expression was observed returning to baseline control by day 70. COMP is a key protein present in cartilage extracellular matrix and is a target of degradation in early OA (30). In addition, proteins such as gelsolin, which had increased expression during the OA+MSC group, has previously been attributed to chondrocyte migration. It could be postulated that proteins highly associated with the joint may be sourced from EVs secreted by joint cells, raising the question of how EVs interact in their *in vivo* environment in response to stimuli.

The most significant GO term associated with DE EV-proteins was serine endopeptidase activity following MSC treatment. Serine type endopeptidases, or serine proteinases have been attributed to proteolytic cartilage destruction. In addition, serine proteinases perform vital functions such as cytokine regulation and receptor activation (31). Degradomic studies have demonstrated that an increase in proteases activity in OA, such as in HtrA1 was responsible for cartilage proteolysis. (32). HtrA1 decreased across time following MSC injection. In a murine models the genetic removal of HtrA1 delayed the degradation of articular or condylar cartilage in mice (33). Moreover, a previous study profiling the synovial fluid-derived EV proteome cargo in OA patients of both sexes identified sex specific differences in cargo, with enriched pathways including proteins involved in endopeptidase activity, specifically in women (34). This is of note as all horses included in this study were mares. With respect to serine endopeptidase activity in MSCs it has previously been reported that interactions between BM-MSCs and natural killer cells are fundamental to improving MSC therapeutic efficacy. It has been stated that serpin B9 has a cytoprotective function in MSCs (35). In other diseases such as colorectal cancer, the use of MSCs identified serpins as having immunomodulatory effects, acting on immune cells in order to induce a wound healing phenotype, as well as angiogenesis and epithelial to mesenchymal transition (36). Therefore, we hypothesize that the change in SF-EV proteome post MSC injection has the capacity to affect the serine endopeptidase pathway that is known for its detrimental effect on cartilage degradation.

A further altered GO term included collagen containing extracellular matrix, often linked with joint homeostasis. Exosomes from embryonic MSCs were found to balance synthesis and degradation of the cartilage extracellular matrix in an *in vitro* murine model (37). In our study, we observe a significant change in the expression of proteins associated with cartilage structure, such as COMP, hyaluronan binding protein 2, cartilage intermediate layer protein, chondroadherin and gelsolin. These proteins return to baseline control level by the end of our study. This suggests a restorative effect and return towards a healthy cartilage phenotype. It may be that such increased expression is more likely to be attributed to EVs secreted by native tissues than MSCs, which contribute to collagen extracellular matrix homeostatic function which could be beneficial in OA treatment. In addition, MSCs could be upstream regulators of these effects from native tissues.

In this study, the level of complement proteins in SF EVs decreased with time and returned to baseline in horses treated with MSCs, potentially affecting disease progression. The complement cascade was also a significantly enriched pathway. This pathway is activated in the early stages of OA, with C3a and C5a attributed to OA progression (38). In addition it has been linked to extracellular cartilage matrix degradation, chondrocyte and synoviocyte inflammatory responses, cell lysis, synovitis, disbalanced bone remodeling, and osteophyte formation (39). Several complement components have previously been identified as upregulated in OA SF. It was reported that C3a and C5a promoted chemotaxis of neutrophils and monocytes, and increased leukotriene synthesis (40). In our study multiple complement factors were identified, including C7, C8, C9 and C2 (C3/C5 convertase) when comparing control to OA+MSCs across time.

There are a number of limitations to our study. The duration of the study only enabled the quantification of the effect of MSCs *in vivo* on the global EV proteome in the short term. MSC viability following treatment could not be quantified. Thus, there is an inability to determine the number of EVs contributing to the SF-EV proteome derived from injected MSCs and those from joint tissues themselves. In addition, it is poorly characterised how long MSCs survive *in vivo* and how it affects their EV production. Various animal models have also been employed to quantify the biodistribution of MSCs. In a murine model it was found that EV source and route of administration affected biodistribution *in vivo* (41). An addition to this experiment would include a group in which OA was induced but not treated with MSCs. The severity of OA phenotype is also a limitation, as the model has been shown to produce a post-traumatic OA phenotype with respect to clinical parameters, but molecularly there were no significant proteins identified between control and OA. Prior to FDR correction ANOVA identified 39 DE proteins with respect to group (control and OA), including: Spondin-1 (p=0.005), HtrA1 serine endopeptidase (p=0.02), and serotransferrin (p=0.01), all of which have been previously implicated in OA pathogenesis (42–44). Thus whilst we cannot be certain there appears to be an effect on the EV proteome following OA induction that would require further validation.

It needs to be determined if the SF were acting in a causative or reactionary manner to the *in vivo* environment. There is a significant degree of variability with respect to MSC properties dependent on the tissue source. Our study used adipose-derived cells and different results may have been achieved with alternative source of MSCs. A recent study by Roelefs *et al*. found that synovium-derived adult GDF5-lineage MSCs had a significant role in response to joint injury, and a higher affinity for cartilage repair (45). Thus, it is likely that the EV cargo will be MSC source dependent, potentially for the MSC-derived EVs as well as the joint tissue-EV response to MSCs.

Ultimately, we hypothesize that after the introduction of MSCs into the joint, MSC-EVs deliver proteinous cargo into recipient cells found within the intra-articular environment, while also promoting intrinsic cellular changes altering the cargo of EVs secreted from native cells. We suggest they act partially through effects on the serine endopeptidase pathway, subsequently reducing its activity and OA pathogenic effect. In addition, altered EV proteins are implicated in the complement system and collagen containing extracellular matrix pathway, as shown in Figure 8. These altered pathways may provide potential targets for therapeutic intervention and require further explorationin the context of MSC therapy and the use of MSC-EVs in OA. It is important to determine if EVs have the same immunogenic effect as allogenic MSCs, and determine the effect of other variables such as sex and tissue type.

**Figure 8.**
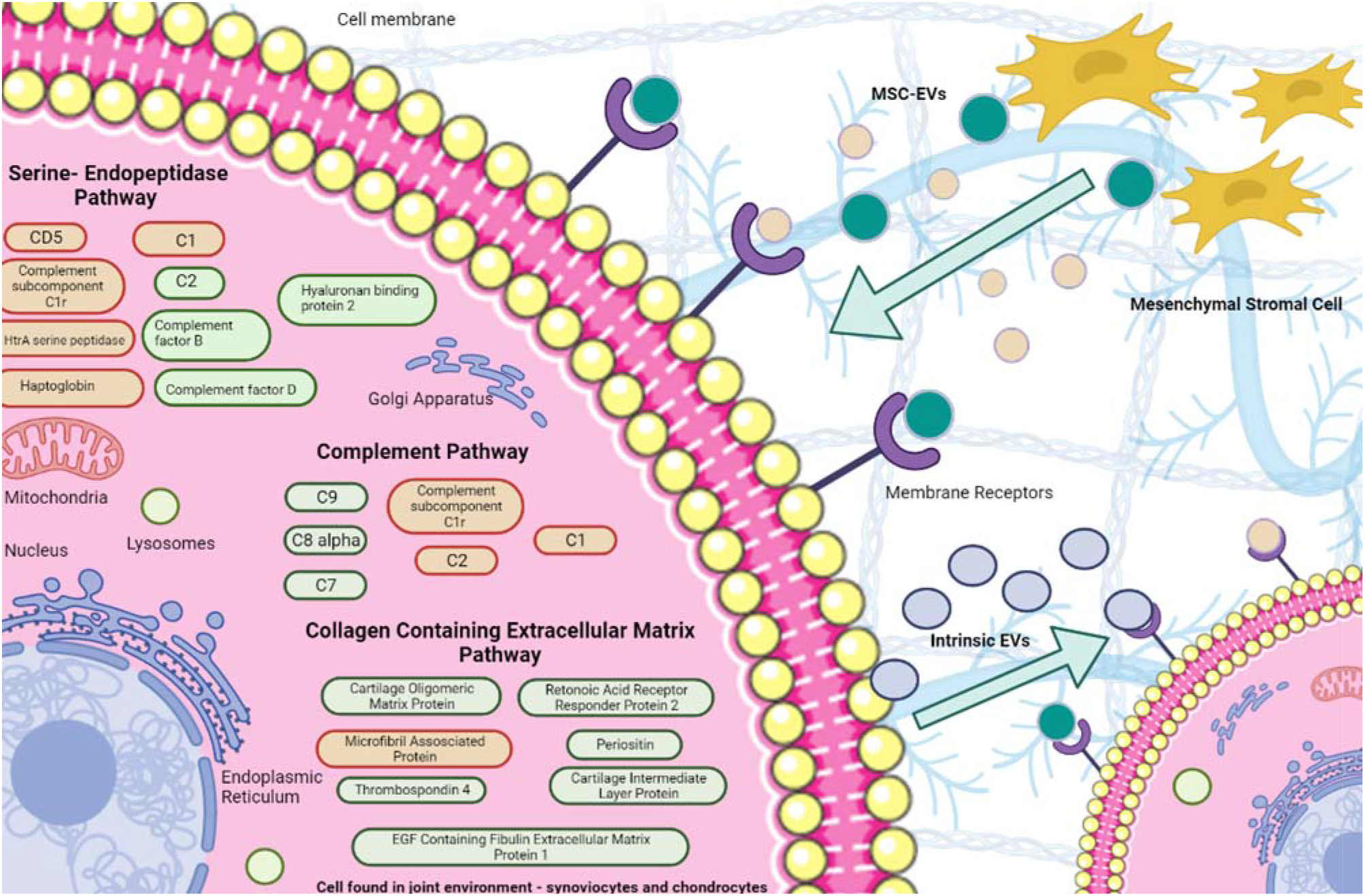
Potential mechanisms of action of SF-EVs following MSC treatment. EVs were sourced from both MSCs and the intraarticular environment. We hypothesized that MSC-EVs affect the intraarticular cells through their differential protein cargo, resulting in altered EV secretion from intrinsic cells. DE proteins attributed to given pathways are red (decreased expression to meet baseline or surpass it by day 70) or green (increased expression to reach baseline at day 70).

## 5 Conclusion

We characterized for the first time using an unbiased approach the SF-EV protein cargo in a model of OA after MSC administration. Changes in the proteome of the synovial fluid-derived EVs following MSC injection are suggestive of EVs playing a role in mediating the effect of cell therapy. A time dependent change in potential therapeutic efficacy of allogenic MSCs was also observed. Potential targets were identified that warrant further investigation in order to determine their significance in pathophysiology and management of equine OA.

## Supporting information

Supplementary figures and tables - word

Tables and Figures word

## 6 Conflicts of Interest

The authors declare that the research was conducted in the absence of any commercial or financial relationships that could be construed as a potential conflict of interest.

## 7 Author Contributions

Conceptualization, MJP, VJ and SJ.; methodology, EJC, MP, SJ, CA, LCB, EJ, ECG, RJ, AT; formal analysis, EJC, EJ, ECG, RJ, AS; investigation, EJC, EJ, RJ.; data curation, EJC,RJ, EJ,AS.; writing—original draft preparation, EJC,; writing—review and editing, EJC, MP, SJ, CA, LB, EJ, ECG, RJ, AS, AT,LCB, CL,KU, ELA; visualization, EJC, RJ, AS, EJ, ECG.; all authors have read and agreed to the published version of the manuscript.

## 8 Funding

Emily Clarke is a self-funded PhD student. Mandy Peffers is funded through a Wellcome Trust Intermediate Clinical Fellowship (107471/Z/15/Z). This work was supported by the Horserace Betting Levy Board in conjunction with the racing foundation (SPrj048). Our work is also supported by the Medical Research Council (MRC) and Versus Arthritis as part of the MRC Versus Arthritis Centre for Integrated research into Musculoskeletal Ageing (CIMA). This work was funded by the Horserace Betting Levy Board (HBLB), project code SPrj048. The Authors acknowledge use of the CDSS Bioanalytical Facility provided by Liverpool Shared Research Facility Voucher (Facilities, Faculty of Health and Life Sciences, University of Liverpool).

## 10 Supplementary Tables and Figures

**Table 1.**
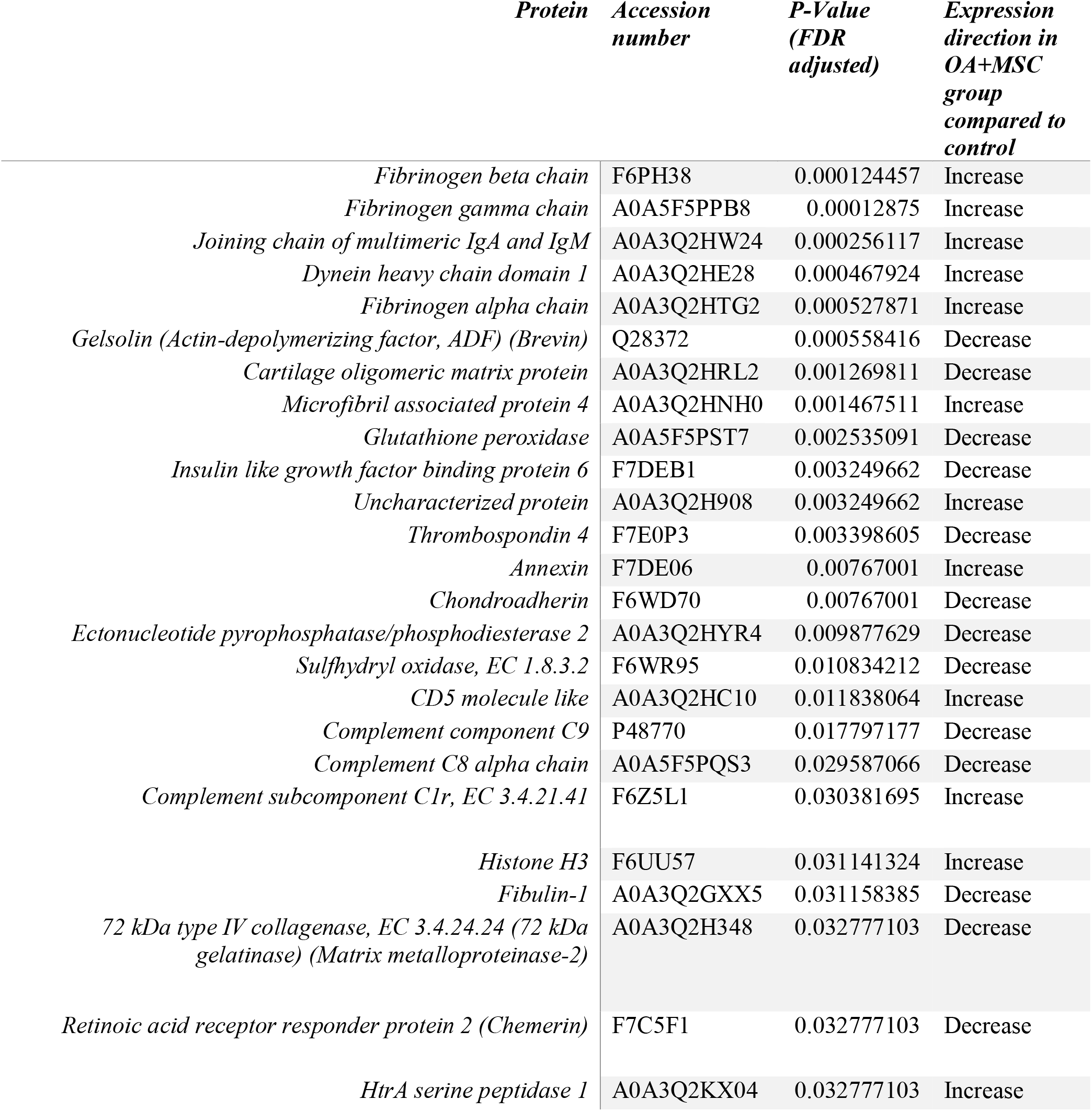
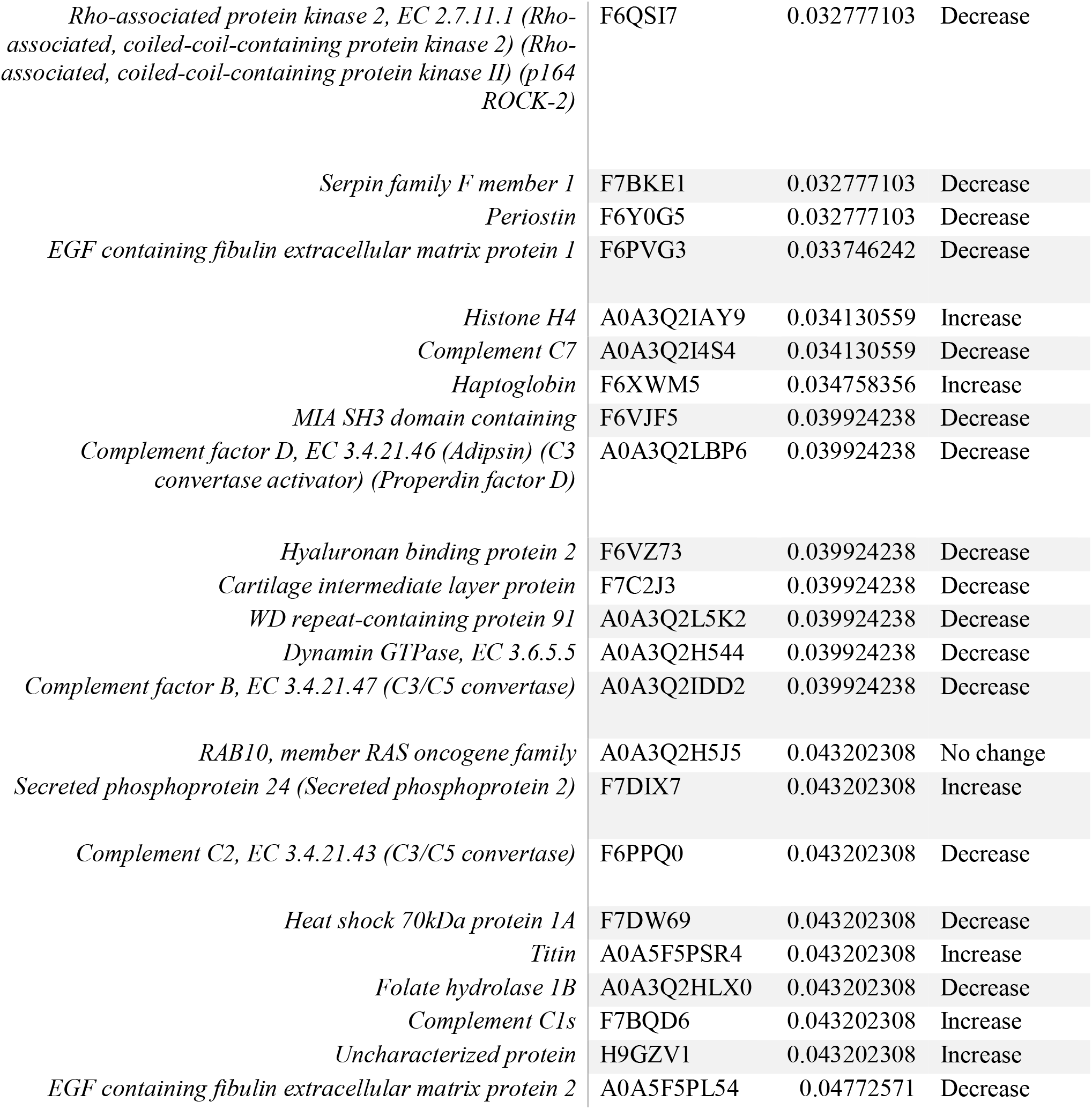
A table showing all 48 significant proteins, with accompanying accession number, FDR corrected P-Value, and direction of expression from MSC injection into the joint, to the end of the 70 day study. All proteins returning to baseline control after 70 days.

**Figure 1.**
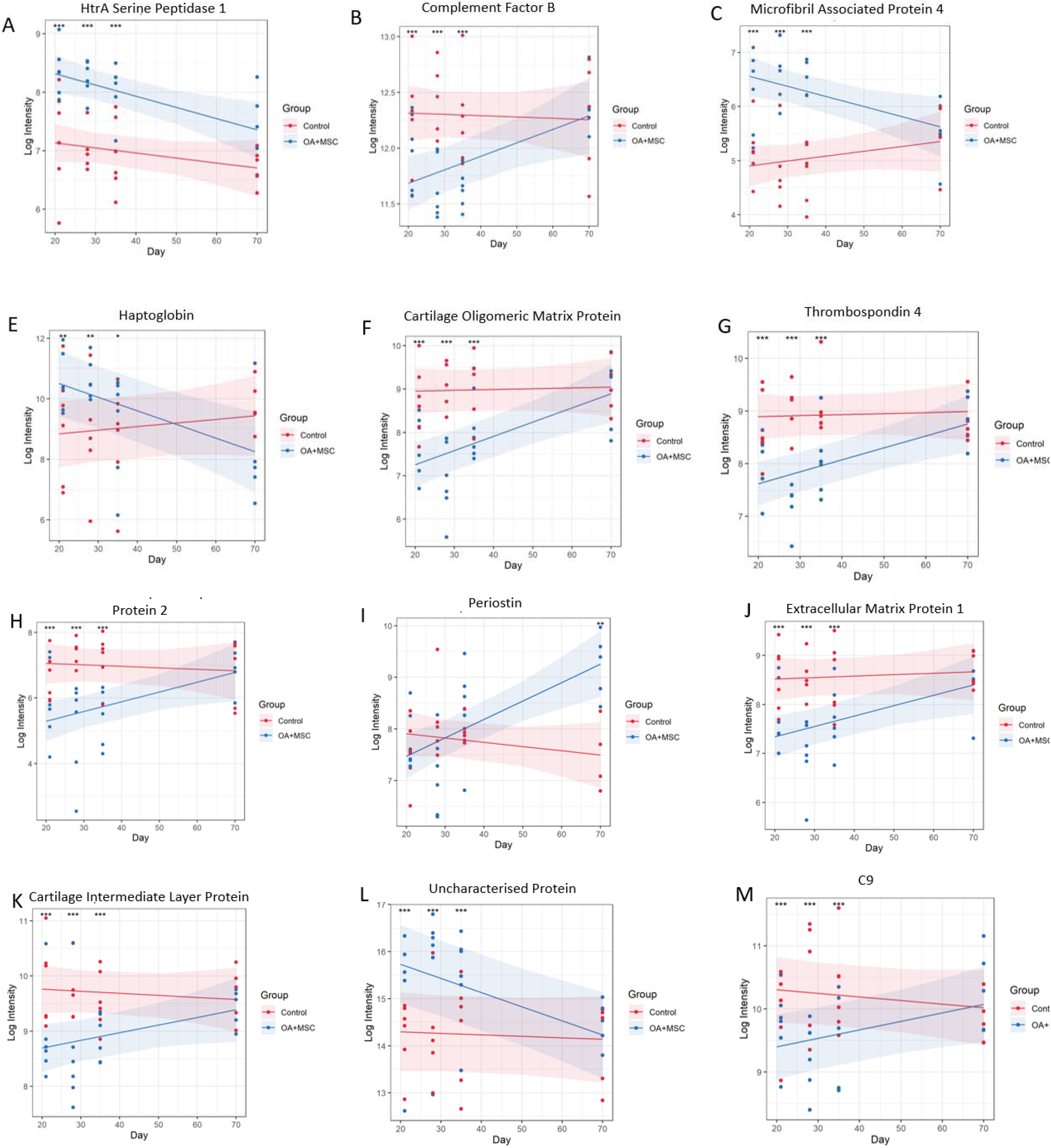
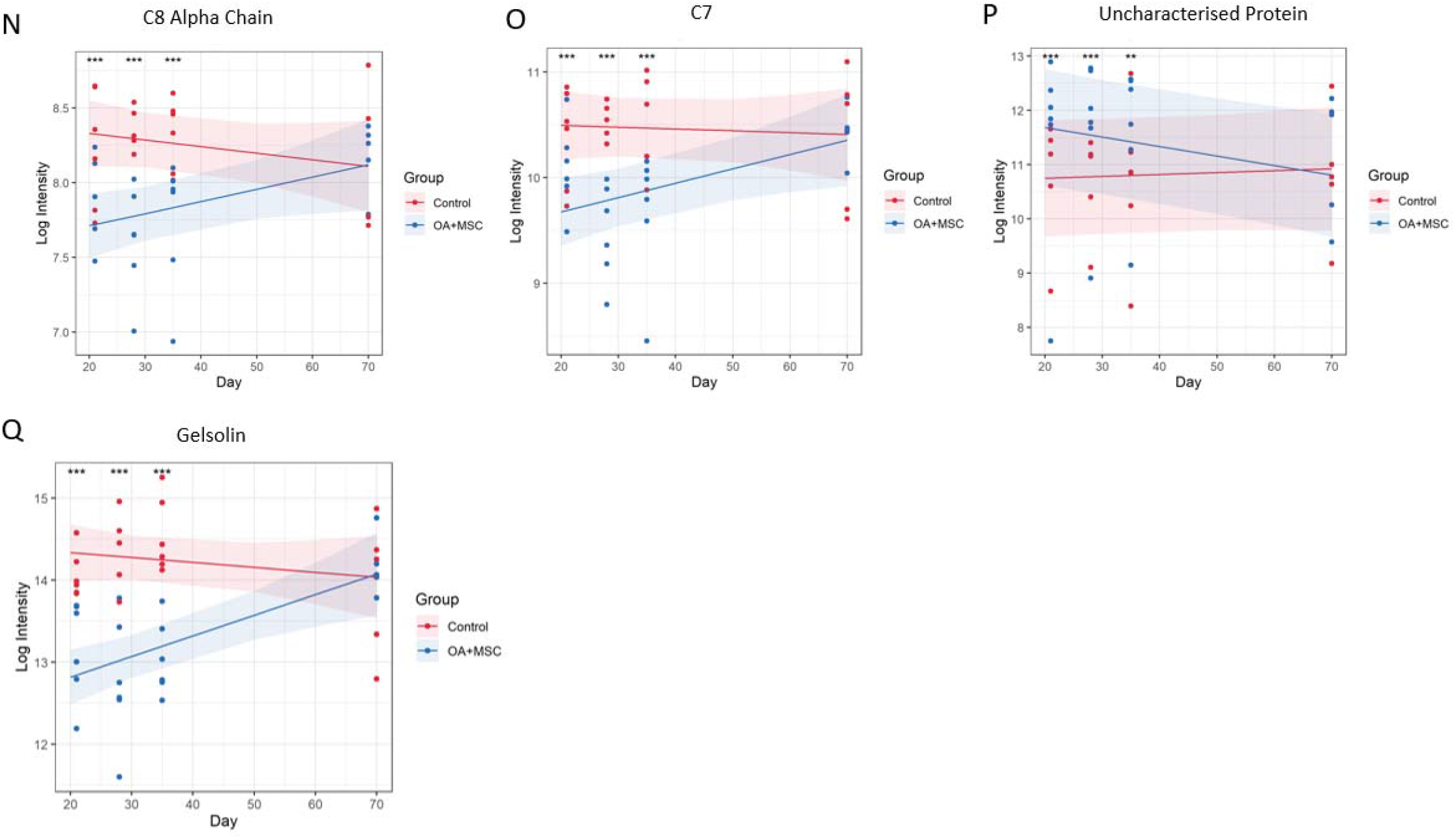
Differentially abundant proteins mapped to one of the three most significant functional enrichment pathway. Graphs demonstrating the results of significant (P≤ 0.05) proteins identified after a linear mixed model was applied to the experimental cohort, comparing protein expression between experimental group (treatment OA+MSCs (treatment) and control over time (day 21, 28, 35 and 70), visualizing the expression change longitudinally. Pairwise comparisons were conducted post linear mixed model application to compare control and OA +MSCs protein expression per time point. Significance level, as determined by the generated FDR corrected P Value is shown using (P<0.05, *; P<0.01 **: P<0.001, ***, P<0.0001, ****).

## Notes

### Competing Interest Statement

The authors have declared no competing interest.

