## Supplementary Figures and Tables for "Temporal Extracellular Vesicle Protein Changes following Intraarticular Treatment with Integrin α10β1-selected Mesenchymal Stem Cells in Equine Osteoarthritis"

### Supplementary Tables and Figures

**Table 1.** A table showing all 48 significant proteins, with accompanying accession number, FDR corrected P-Value, and direction of expression from MSC injection into the joint, to the end of the 70 day study. All proteins returning to baseline control after 70 days.

| *Protein* | *Accession number* | *P-Value (FDR adjusted)* | *Expression direction in OA+MSC group compared to control* |
|---|---|---|---|
| *Fibrinogen beta chain* | F6PH38 | 0.000124457 | Increase |
| *Fibrinogen gamma chain* | A0A5F5PPB8 | 0.00012875 | Increase |
| *Joining chain of multimeric IgA and IgM* | A0A3Q2HW24 | 0.000256117 | Increase |
| *Dynein heavy chain domain 1* | A0A3Q2HE28 | 0.000467924 | Increase |
| *Fibrinogen alpha chain* | A0A3Q2HTG2 | 0.000527871 | Increase |
| *Gelsolin (Actin-depolymerizing factor, ADF) (Brevin)* | Q28372 | 0.000558416 | Decrease |
| *Cartilage oligomeric matrix protein* | A0A3Q2HRL2 | 0.001269811 | Decrease |
| *Microfibril associated protein 4* | A0A3Q2HNH0 | 0.001467511 | Increase |
| *Glutathione peroxidase* | A0A5F5PST7 | 0.002535091 | Decrease |
| *Insulin like growth factor binding protein 6* | F7DEB1 | 0.003249662 | Decrease |
| *Uncharacterized protein* | A0A3Q2H908 | 0.003249662 | Increase |
| *Thrombospondin 4* | F7E0P3 | 0.003398605 | Decrease |
| *Annexin* | F7DE06 | 0.00767001 | Increase |
| *Chondroadherin* | F6WD70 | 0.00767001 | Decrease |
| *Ectonucleotide pyrophosphatase/phosphodiesterase 2* | A0A3Q2HYR4 | 0.009877629 | Decrease |
| *Sulfhydryl oxidase, EC 1.8.3.2* | F6WR95 | 0.010834212 | Decrease |
| *CD5 molecule like* | A0A3Q2HC10 | 0.011838064 | Increase |
| *Complement component C9* | P48770 | 0.017797177 | Decrease |

| | | | |
|---|---|---|---|
| *Complement C8 alpha chain* | A0A5F5PQS3 | 0.029587066 | Decrease |
| *Complement subcomponent C1r, EC 3.4.21.41* | F6Z5L1 | 0.030381695 | Increase |
| *Histone H3* | F6UU57 | 0.031141324 | Increase |
| *Fibulin-1* | A0A3Q2GXX5 | 0.031158385 | Decrease |
| *72 kDa type IV collagenase, EC 3.4.24.24 (72 kDa gelatinase) (Matrix metalloproteinase-2)* | A0A3Q2H348 | 0.032777103 | Decrease |
| *Retinoic acid receptor responder protein 2 (Chemerin)* | F7C5F1 | 0.032777103 | Decrease |
| *HtrA serine peptidase 1* | A0A3Q2KX04 | 0.032777103 | Increase |
| *Rho-associated protein kinase 2, EC 2.7.11.1 (Rho-associated, coiled-coil-containing protein kinase 2) (Rho-associated, coiled-coil-containing protein kinase II) (p164 ROCK-2)* | F6QSI7 | 0.032777103 | Decrease |
| *Serpin family F member 1* | F7BKE1 | 0.032777103 | Decrease |
| *Periostin* | F6Y0G5 | 0.032777103 | Decrease |
| *EGF containing fibulin extracellular matrix protein 1* | F6PVG3 | 0.033746242 | Decrease |
| *Histone H4* | A0A3Q2IAY9 | 0.034130559 | Increase |
| *Complement C7* | A0A3Q2I4S4 | 0.034130559 | Decrease |
| *Haptoglobin* | F6XWM5 | 0.034758356 | Increase |
| *MIA SH3 domain containing* | F6VJF5 | 0.039924238 | Decrease |
| *Complement factor D, EC 3.4.21.46 (Adipsin) (C3 convertase activator) (Properdin factor D)* | A0A3Q2LBP6 | 0.039924238 | Decrease |
| *Hyaluronan binding protein 2* | F6VZ73 | 0.039924238 | Decrease |
| *Cartilage intermediate layer protein* | F7C2J3 | 0.039924238 | Decrease |
| *WD repeat-containing protein 91* | A0A3Q2L5K2 | 0.039924238 | Decrease |
| *Dynamin GTPase, EC 3.6.5.5* | A0A3Q2H544 | 0.039924238 | Decrease |
| *Complement factor B, EC 3.4.21.47 (C3/C5 convertase)* | A0A3Q2IDD2 | 0.039924238 | Decrease |
| *RAB10, member RAS oncogene family* | A0A3Q2H5J5 | 0.043202308 | No change |
| *Secreted phosphoprotein 24 (Secreted phosphoprotein 2)* | F7DIX7 | 0.043202308 | Increase |

| | | | |
|---|---|---|---|
| *Complement C2, EC 3.4.21.43 (C3/C5 convertase)* | F6PPQ0 | 0.043202308 | Decrease |
| *Heat shock 70kDa protein 1A* | F7DW69 | 0.043202308 | Decrease |
| *Titin* | A0A5F5PSR4 | 0.043202308 | Increase |
| *Folate hydrolase 1B* | A0A3Q2HLX0 | 0.043202308 | Decrease |
| *Complement C1s* | F7BQD6 | 0.043202308 | Increase |
| *Uncharacterized protein* | H9GZV1 | 0.043202308 | Increase |
| *EGF containing fibulin extracellular matrix protein 2* | A0A5F5PL54 | 0.04772571 | Decrease |

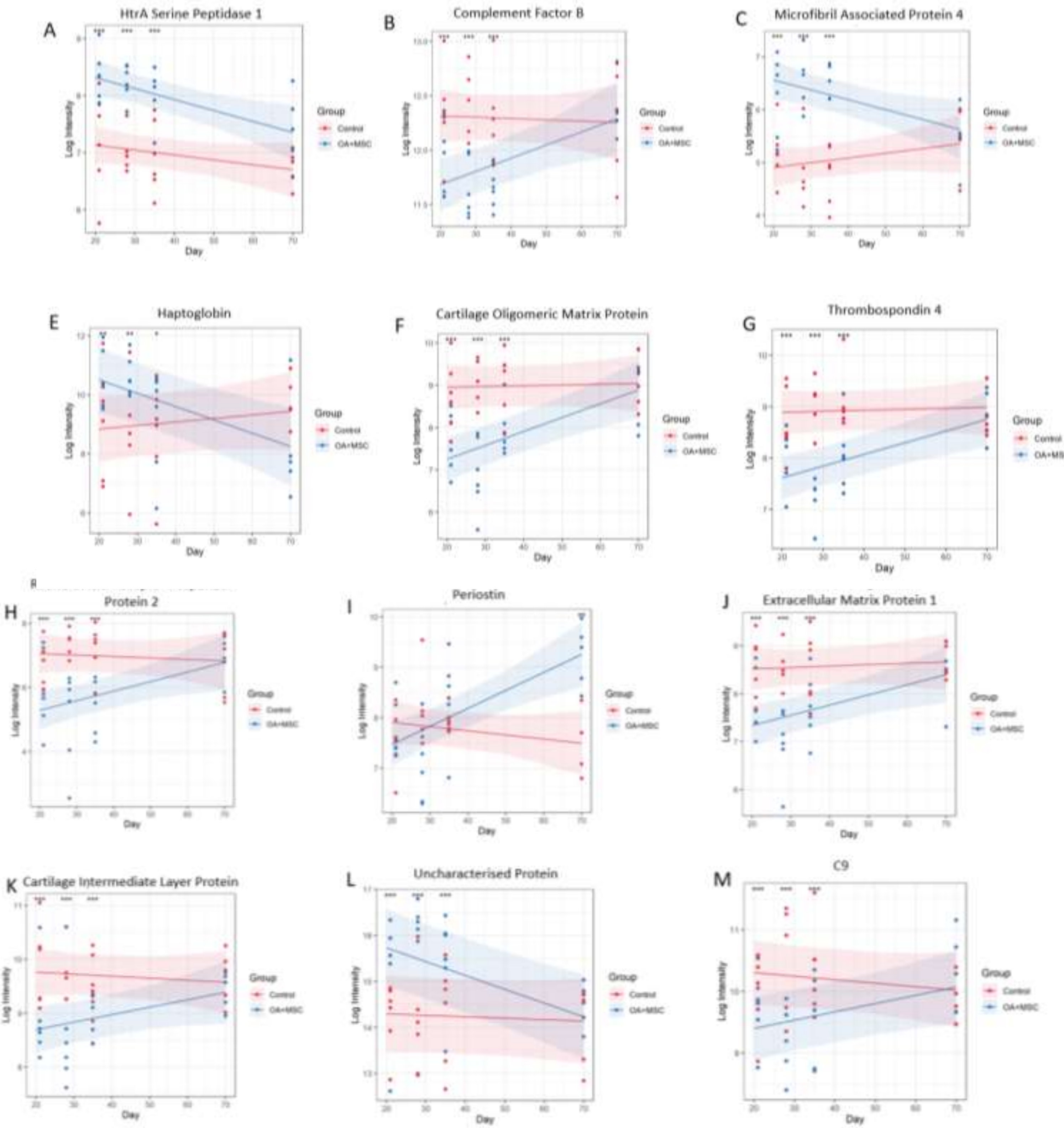
A
HtrA Serine Peptidase 1
B
Complement Factor B
C
Microfibril Associated Protein 4
E
Haptoglobin
F
Cartilage Oligomeric Matrix Protein
G
Thrombospondin 4
H
Protein 2
I
Periostin
J
Extracellular Matrix Protein 1
K
Cartilage Intermediate Layer Protein
L
Uncharacterised Protein
M
C9
Group
Control
OA+MSC
Day
Log Intensity

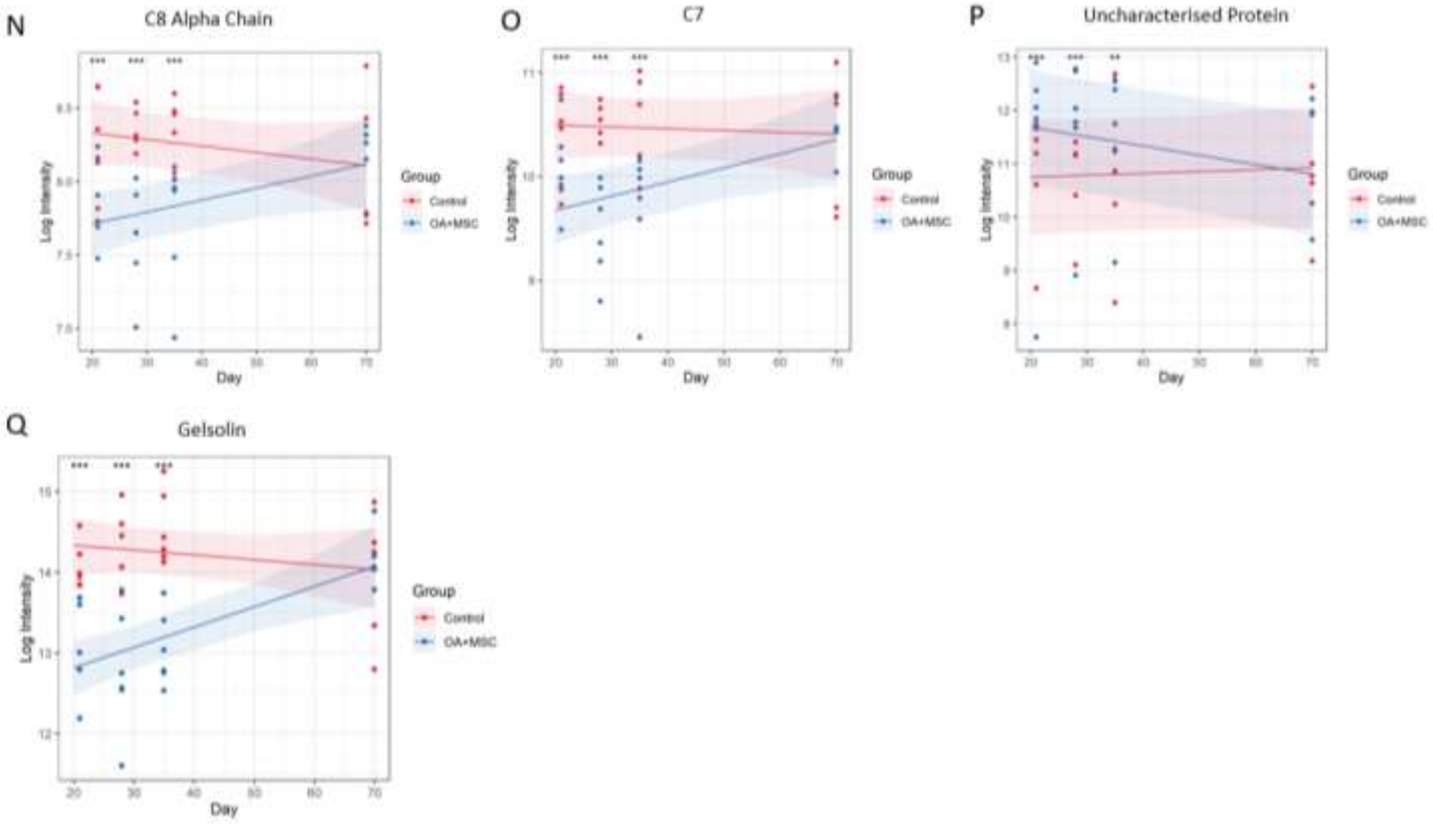


**Figure 1.** Differentially abundant proteins mapped to one of the three most significant functional enrichment pathway. Graphs demonstrating the results of significant ($P \leq 0.05$) proteins identified after a linear mixed model was applied to the experimental cohort, comparing protein expression between experimental group (treatment OA+MSCs (treatment) and control over time (day 21, 28, 35 and 70), visualizing the expression change longitudinally. Pairwise comparisons were conducted post linear mixed model application to compare control and OA +MSCs protein expression per time point. Significance level, as determined by the generated FDR corrected P Value is shown using ($P<0.05$, *; $P<0.01$ **: $P<0.001$, ***, $P<0.0001$, ****).
