## Supplementary material for "Temporal Extracellular Vesicle Protein Changes following Intraarticular Treatment with Integrin α10β1-selected Mesenchymal Stem Cells in Equine Osteoarthritis": Figures and Tables

*Tables and Figures*

**Table 1-.An overview of experimental groups and the longitudinal time points for SF collection.**

| Time point (day) | Control | OA | OA+MSC |
|---|---|---|---|
| 0 (prior to surgery) | Yes | Yes | |
| 18 | Yes | Yes | |
| 21 | Yes | | Yes |
| 28 | Yes | | Yes |
| 35 | Yes | | Yes |
| 70 | Yes | | Yes |

*Abbreviations: Osteoarthritis and mesenchymal stromal cells (OA+MSCs), Osteoarthritis (OA)*

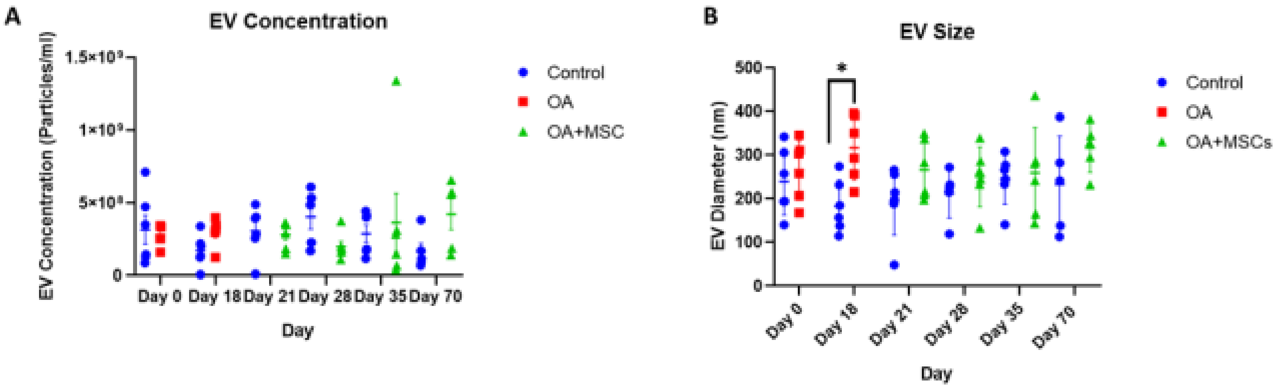


**Figure 1.** Size and concentration of synovial fluid-derived nanoparticles. Nanoparticle tracking analysis was undertaken using a Nanosight NS3000. All error bars are standard error of the mean (SEM). Statistical analysis undertaken in GraphPad Prism 9.0 using Kruskal Wallis Tests with FDR correction and Mann Whitney Tests within time points. (P<0.05, *; P<0.01 **: p<0.001, ***, p<0.0001, ****). (1A) EV concentration and (1B) EV size.

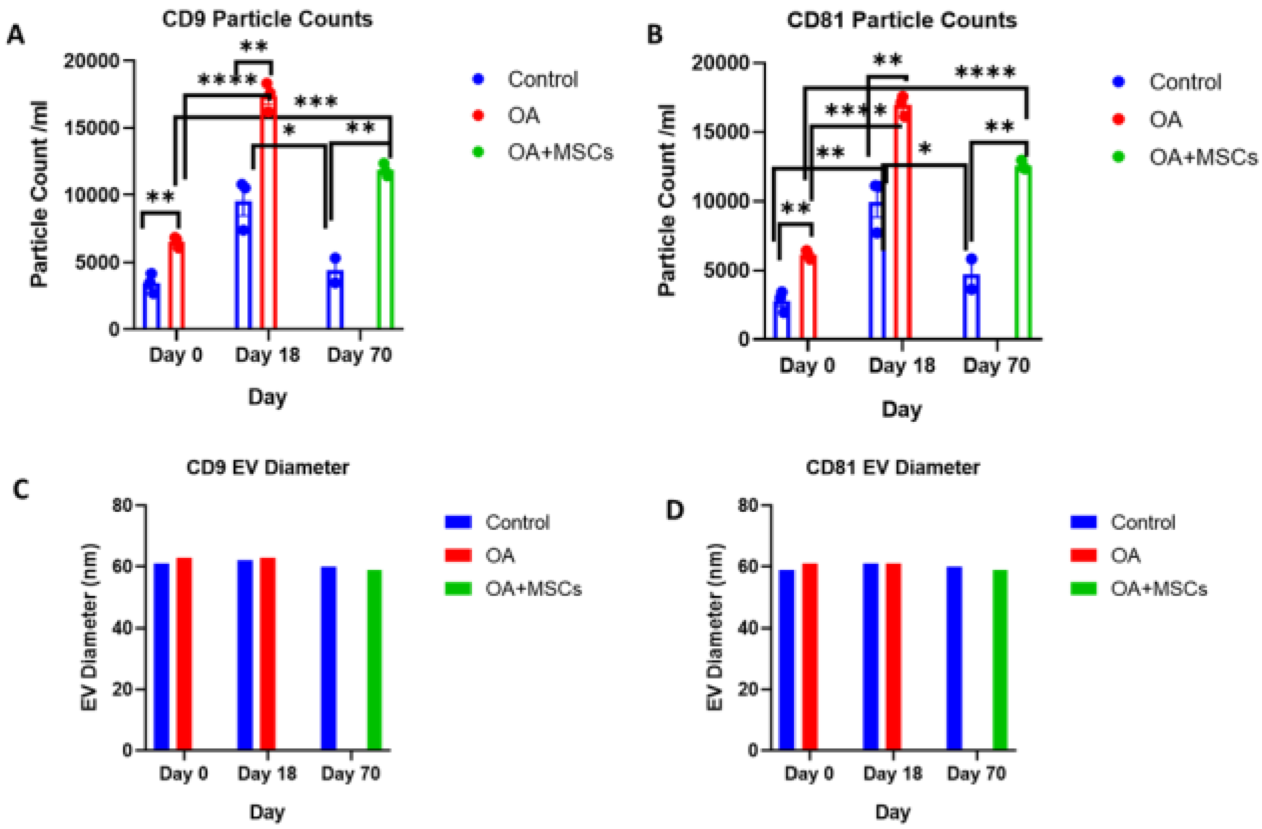


**Figure 2.** Sizing and enumeration of synovial fluid-derived EVs. All data was adjusted for dilution of the sample onto the chip. Shown is the average representing mean of three technical replicates that were run for each sample. Particle numbers were quantified by the number of particles in a defined area on the antibody capture spot. All bars are mean and standard error mean. A. CD9, and B. CD81-positive particles following probing with fluorescent tetraspanin antibodies. C. Sizing of CD9 and D. CD81 labelled EVs, normalised to MIgG control. Limit of detection was 50-200 nm. Statistical analysis undertaken in GraphPad Prism 9.0 using T-tests following parametric evaluation (P<0.05, *; P<0.01 **: P<0.001, ***, P<0.0001, ****).

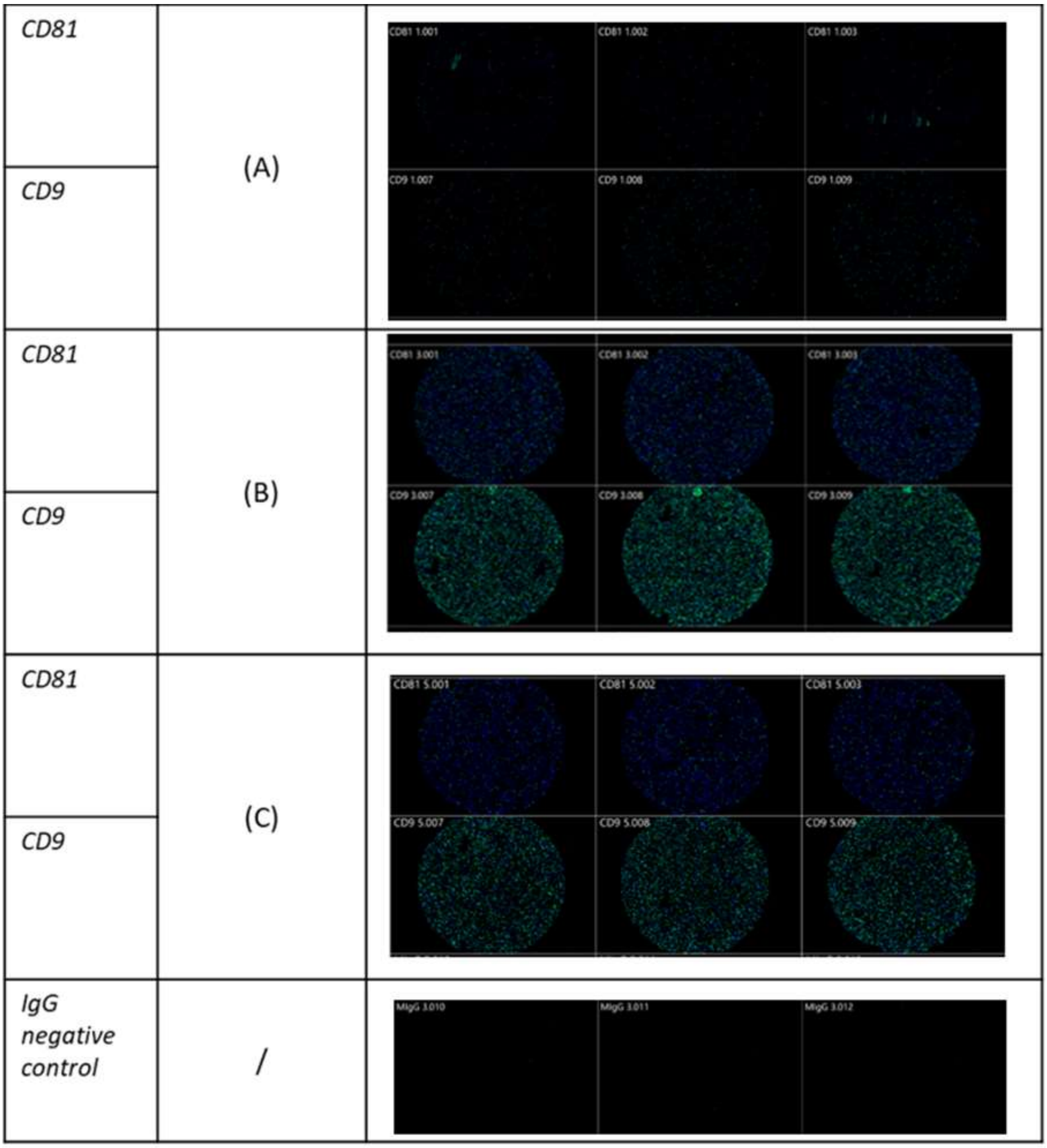


**Figure 3**. Visualization of SF-derived EVs from control, osteoarthritic (OA) and OA +MSCs joints using Exoview at selected time points. A fluorescent image of a representative spot is shown for each sample comparing (A) control, (B) OA and (C) OA +MSCs with colour denoting surface tetraspanin positive identification (blue- CD9, and green -CD81).

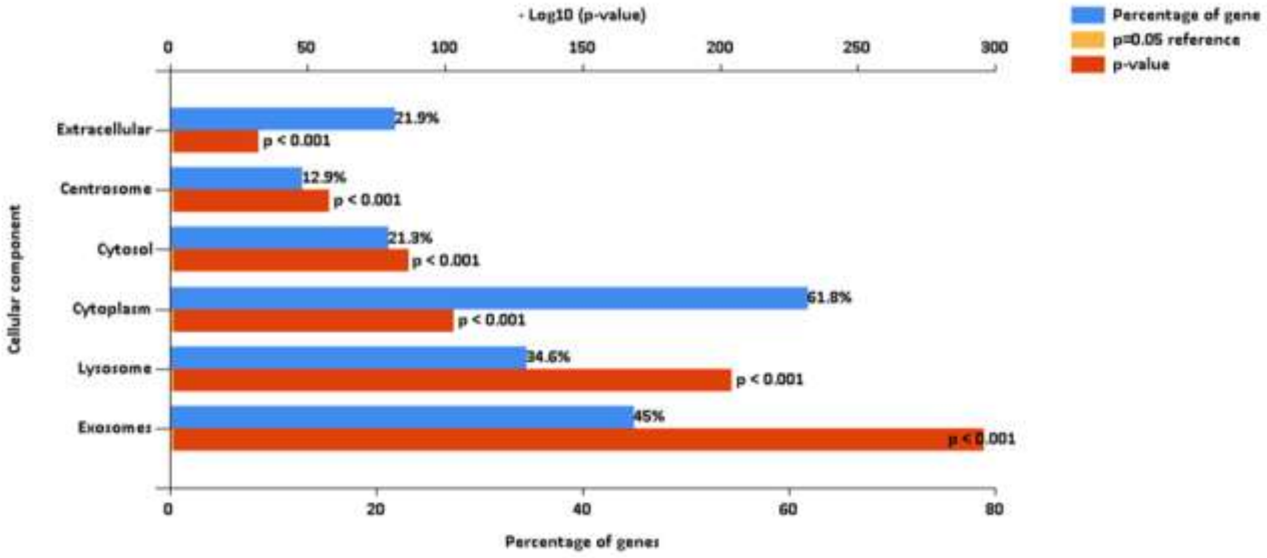


**Figure 4.** FunRich analysis output, after mapping 2047 proteins to GO Cellular Component terms. These proteins were identified from a pooled sample of equine synovial fluid (11ml) used to generate the SF-EV spectral library for this study. SF was sourced from healthy, OA and OA+MSC treated joints in order to encapsulate all potential proteins that may be present across all experimental groups.

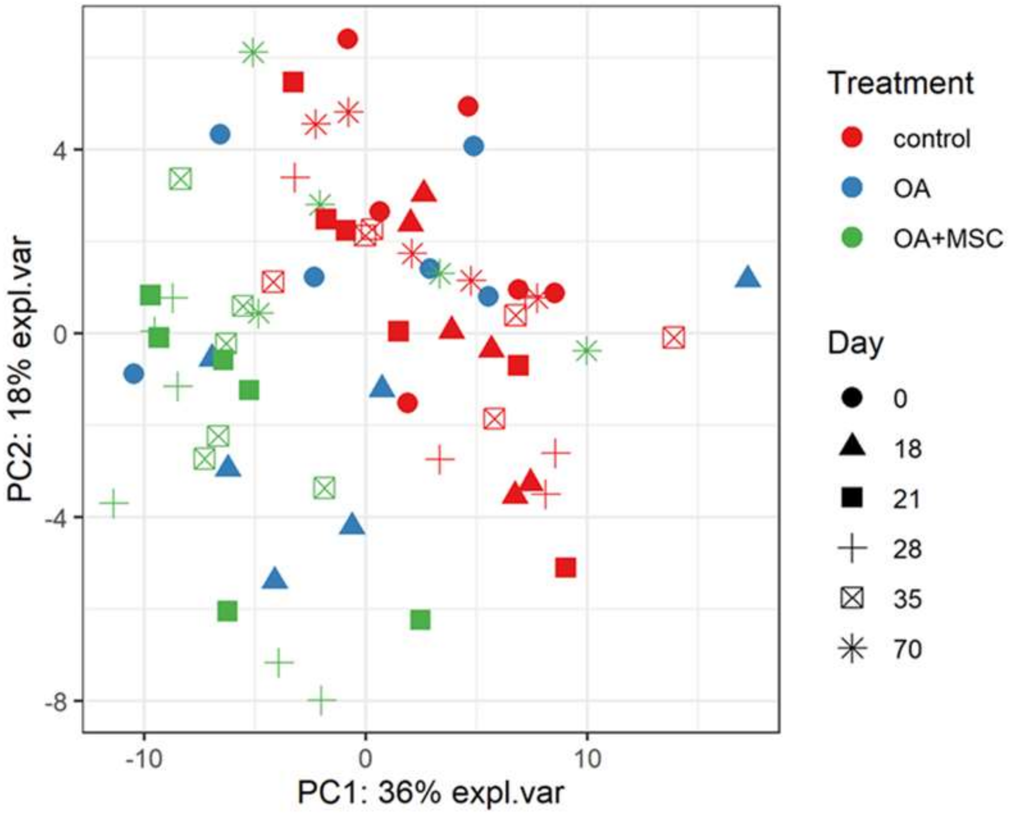


**Figure 5.** Multi-level PCA (mPCA). The first two principal components are plotted, accounting for ~54% of variance. samples based on SWATH-MS. Each plotted point represents a horse, which are coloured by their treatment and shaped by the day of the study.

**Table 2**. The top 10 differentially expressed (P<0.05) proteins accounting for treatment and timepoint. Treatment and time were included as main effects, along with a treatment-by-time interaction term.

| Protein | Accession number | P-Value (FDR adjusted) | Expression direction in OA+MSC group compared to control |
|---|---|---|---|
| **Fibrinogen beta chain** | F6PH38 | 0.0001 | Increase |
| **Fibrinogen gamma chain** | A0A5F5PPB8 | 0.0001 | Increase |
| **Joining chain of multimeric IgA and IgM** | A0A3Q2HW24 | 0.0003 | Increase |
| **Dynein heavy chain domain 1** | A0A3Q2HE28 | 0.0005 | Increase |
| **Fibrinogen alpha chain** | A0A3Q2HTG2 | 0.0005 | Increase |
| **Gelsolin (Actin-depolymerizing factor, ADF) (Brevin)** | Q28372 | 0.0006 | Decrease |
| **Cartilage oligomeric matrix protein** | A0A3Q2HRL2 | 0.001 | Decrease |
| **Microfibril associated protein 4** | A0A3Q2HNH0 | 0.001 | Increase |
| **Glutathione peroxidase** | A0A5F5PST7 | 0.003 | Decrease |
| **Insulin like growth factor binding protein 6** | F7DEB1 | 0.003 | Decrease |

*Abbreviations: Osteoarthritis and mesenchymal stromal cells (OA+MSCs), Immunoglobulin A (IgA) and Immunoglobulin M (IgM)*

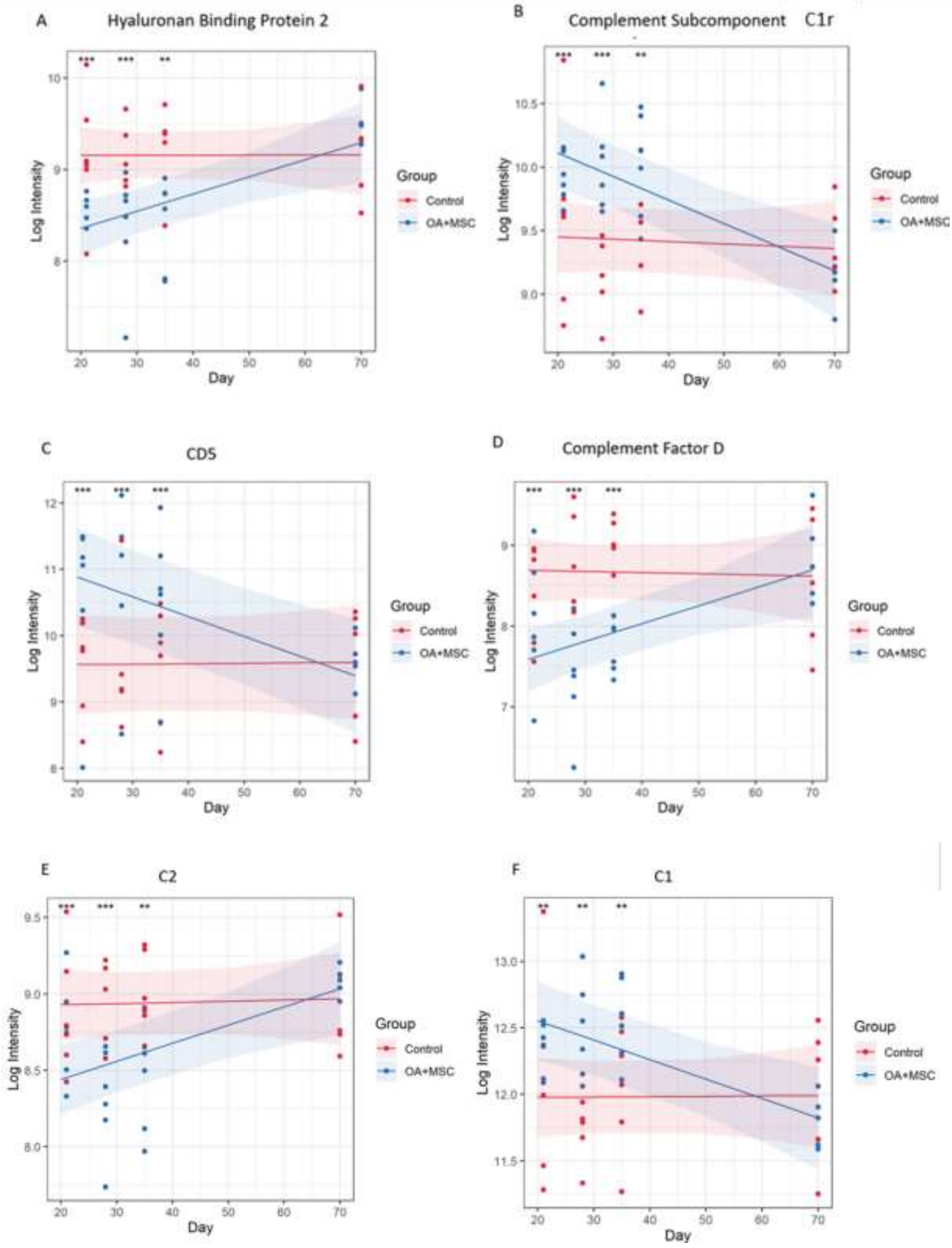


**Figure 6.** Protein expression changes in response to MSC treatment in OA joints vs control. (A) Hyaluronan binding protein 2, (B) Complement subcomponent C1r, (C) CD5, (D) Complement factor D, (E) C2, (F) C1. The models were fitted using the lmerTest implementation of lme4. 48 proteins had a significant group, time, or group:time effect after FDR correction. For each protein with a significant effect the model was plotted using the effects and ggplot2 packages. The fitted model is shown as a line for the OA+MSC (blue) and

control (red). The 95% confidence intervals for each group are shown as a shaded area. The raw data is included as points. At each time point pairwise comparisons were carried out between the treatment and control using the emmeans package. Significance thresholds were as follows: (P<0.05, *; P<0.01 **: P<0.001, ***, P<0.0001, ****).

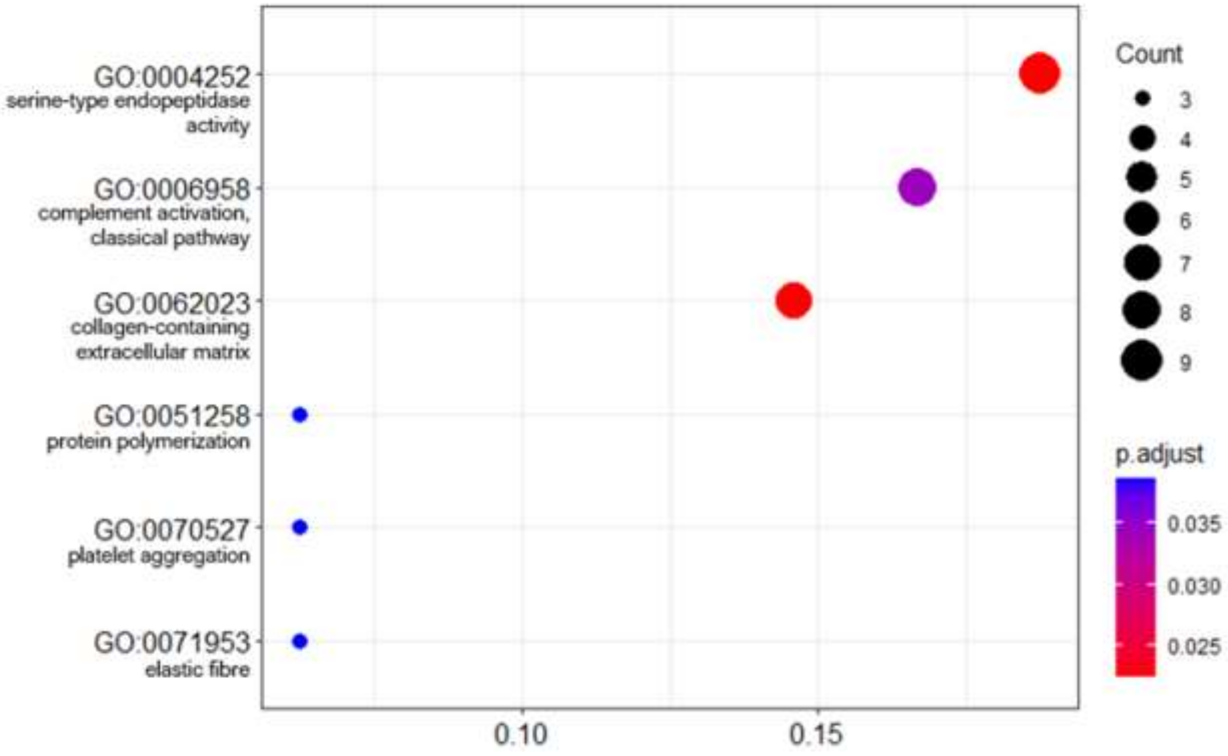


**Figure 7.** Dot plot of GO term enrichment analysis of differentially abundant proteins. The size of the dots indicates the number of proteins that mapped to that term. The x-axis is the protein ratio (number of proteins that map to the term divided by the total number of significant proteins). The dots are shaded by adjusted p-values (BH method).

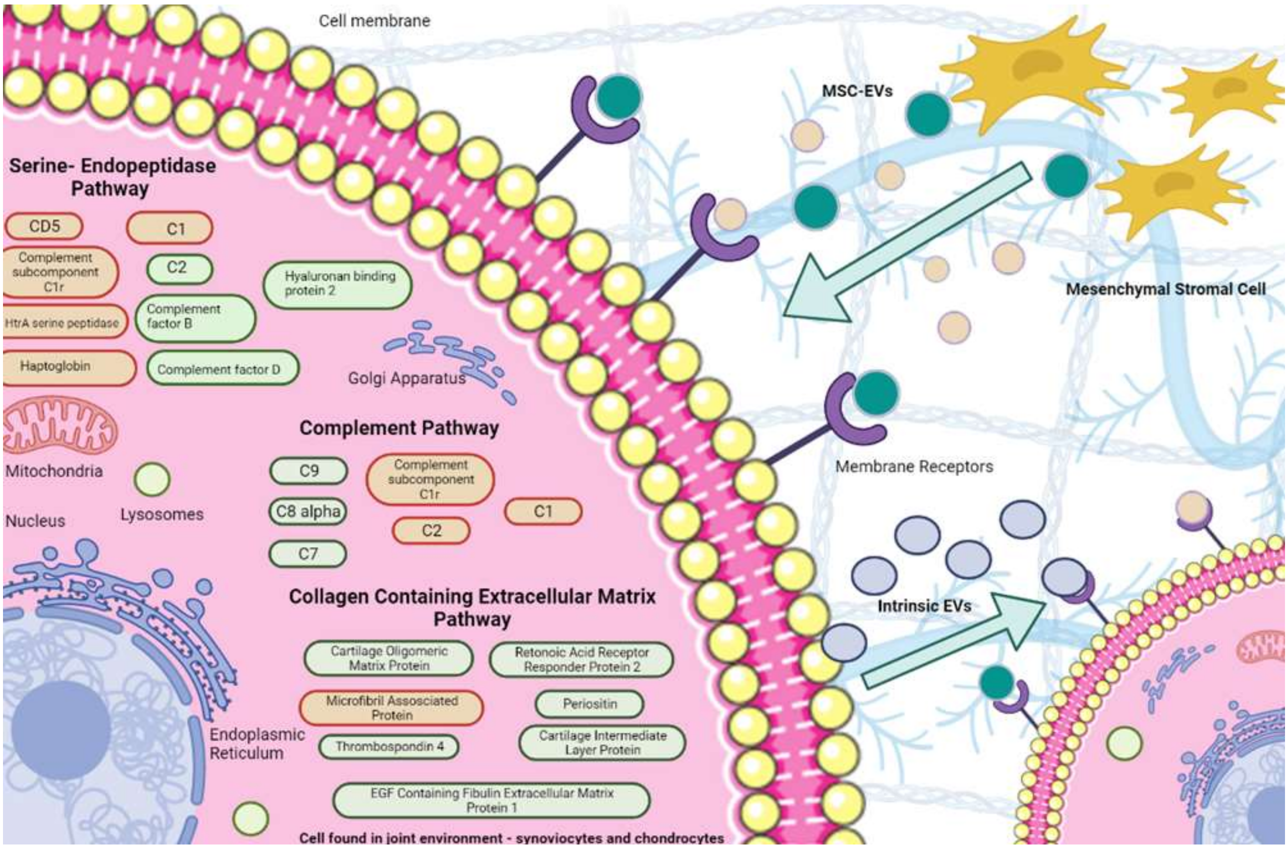


**Figure 8.** Potential mechanisms of action of SF-EVs following MSC treatment. EVs were sourced from both MSCs and the intraarticular environment. We hypothesized that MSC-EVs affect the intraarticular cells through their differential protein cargo, resulting in altered EV secretion from intrinsic cells. DE proteins attributed to given pathways are red (decreased expression to meet baseline or surpass it by day 70) or green (increased expression to reach baseline at day 70).

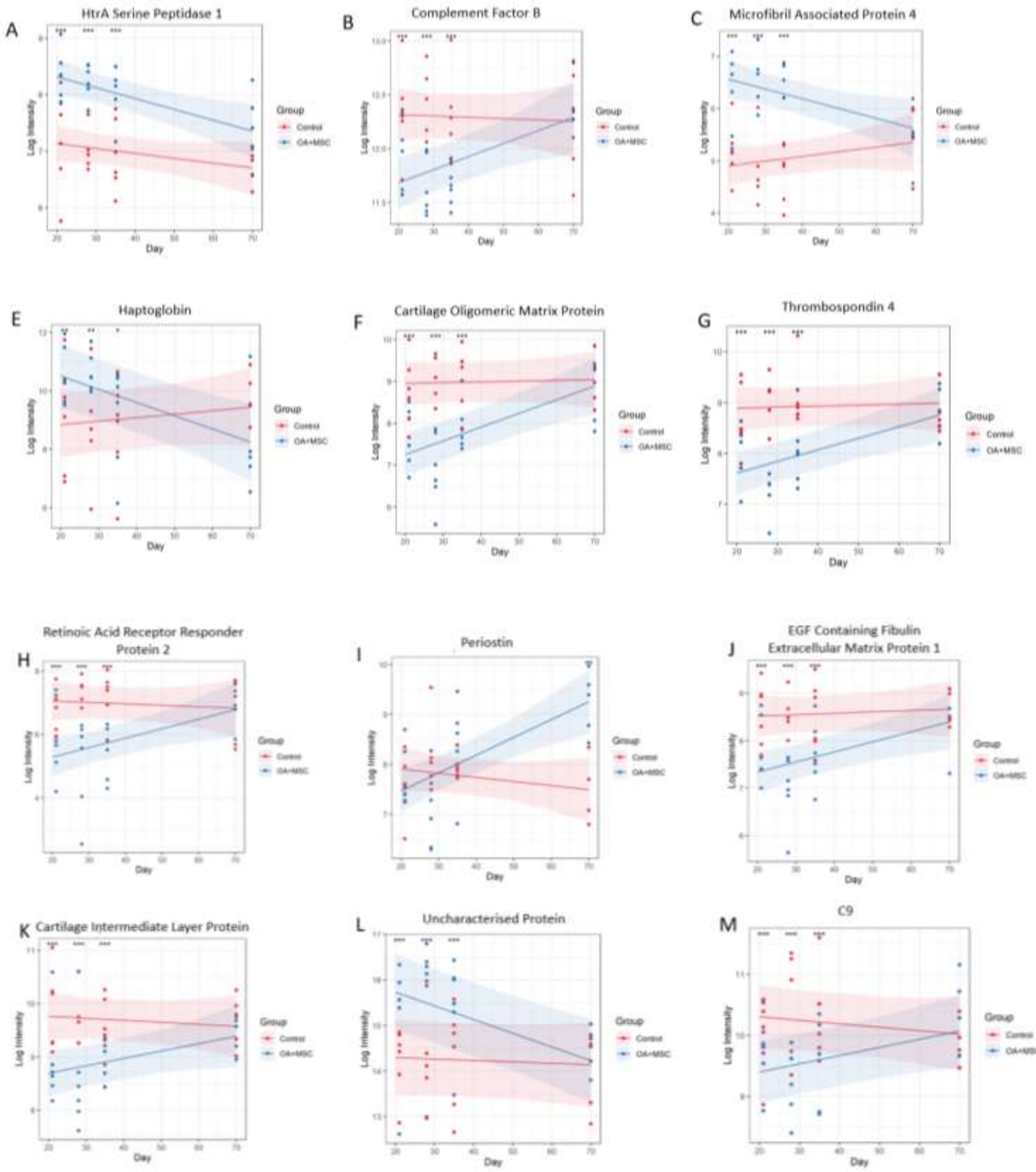

A
HtrA Serine Peptidase 1
B
Complement Factor B
C
Microfibril Associated Protein 4
E
Haptoglobin
F
Cartilage Oligomeric Matrix Protein
G
Thrombospondin 4
H
Retinoic Acid Receptor Responder Protein 2
I
Periostin
J
EGF Containing Fibulin Extracellular Matrix Protein 1
K
Cartilage Intermediate Layer Protein
L
Uncharacterised Protein
M
C9
Group
Control
OA+MSC
Log Intensity
Day

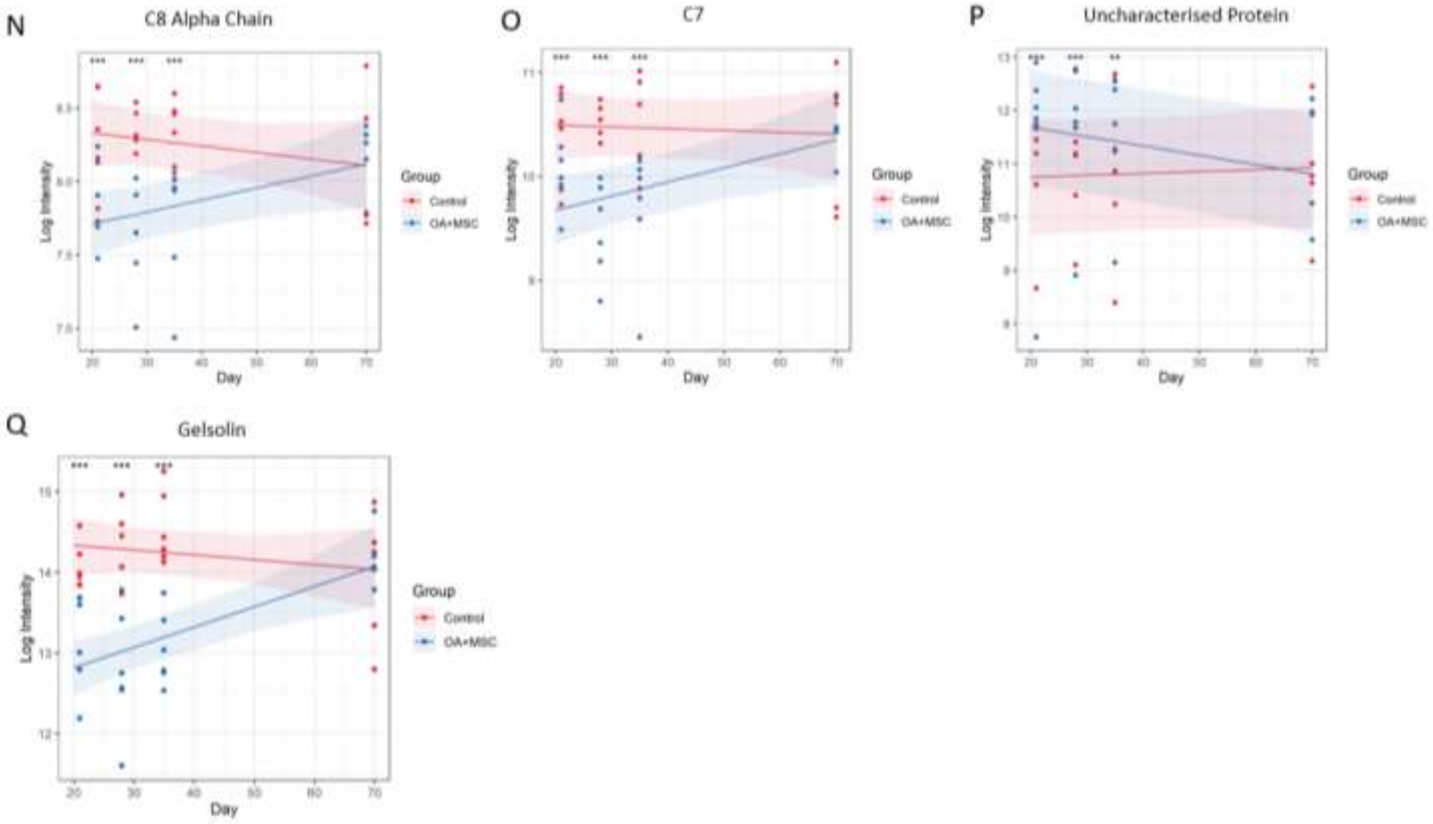


**Figure 1.** Differentially abundant proteins mapped to one of the three most significant functional enrichment pathway. Graphs demonstrating the results of significant (P≤ 0.05) proteins identified after a linear mixed model was applied to the experimental cohort, comparing protein expression between experimental group (treatment OA+MSCs (treatment) and control over time (day 21, 28, 35 and 70), visualizing the expression change longitudinally. Pairwise comparisons were conducted post linear mixed model application to compare control and OA +MSCs protein expression per time point. Significance level, as determined by the generated FDR corrected P Value is shown using (P<0.05, *; P<0.01 **: P<0.001, ***, P<0.0001, ****).
