## Supplementary figures and tables - word for "Temporal Extracellular Vesicle Protein Changes following Intraarticular Treatment with Integrin α10β1-selected Mesenchymal Stem Cells in Equine Osteoarthritis"

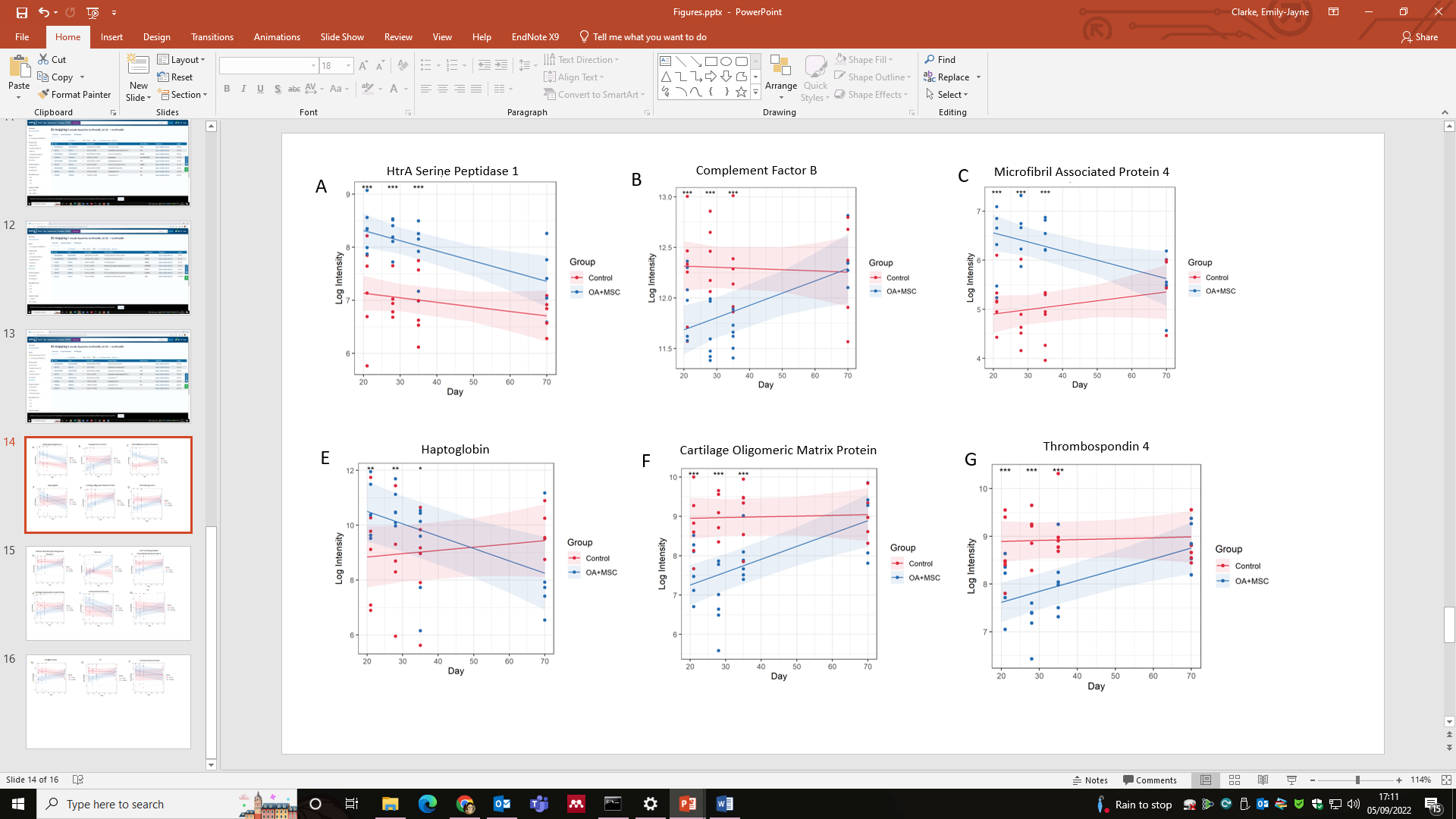

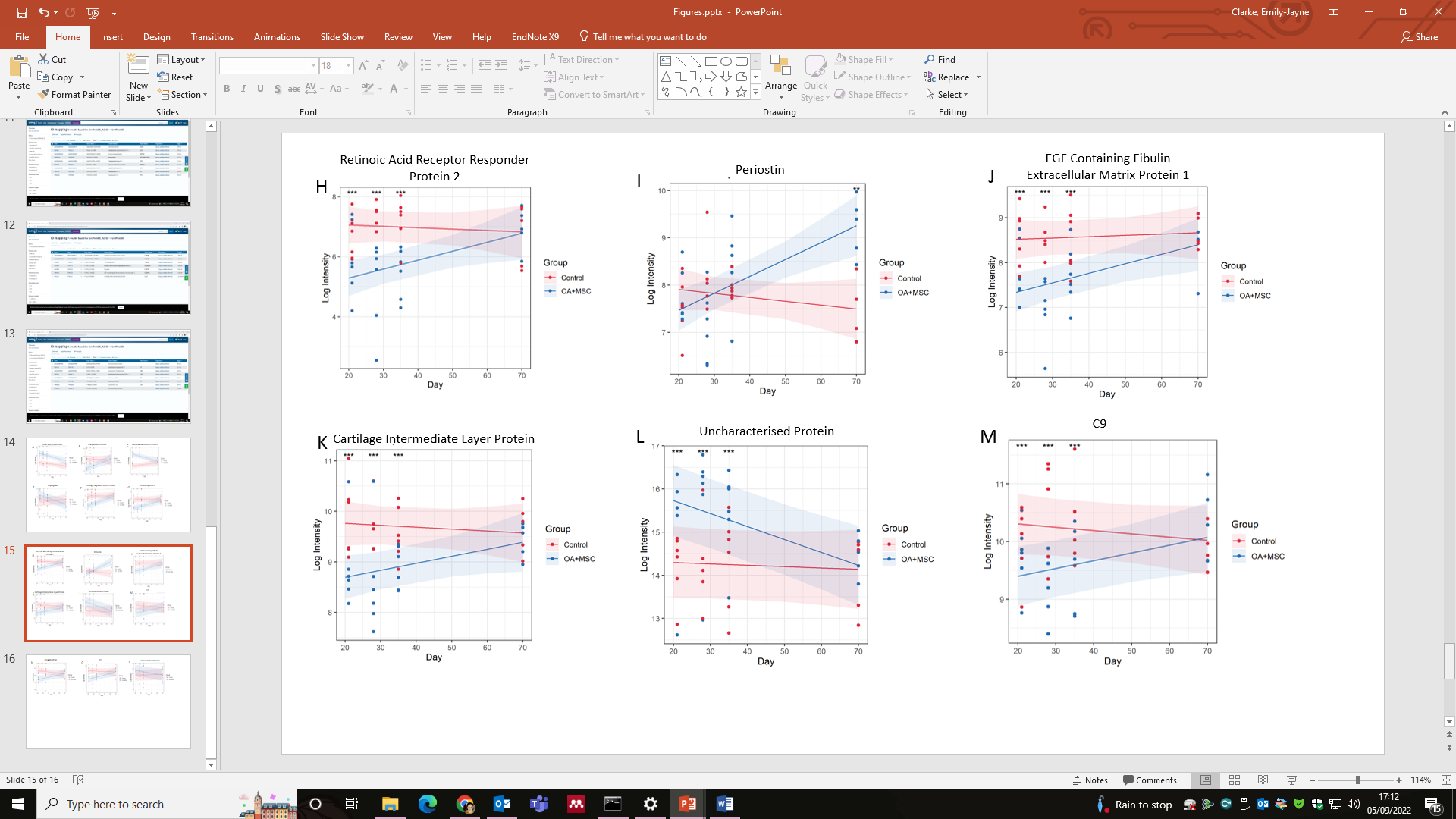

**Figure 1.** Differentially abundant proteins mapped to one of the three most significant functional enrichment pathway. Graphs demonstrating the results of significant (P≤ 0.05) proteins identified after a linear mixed model was applied to the experimental cohort, comparing protein expression
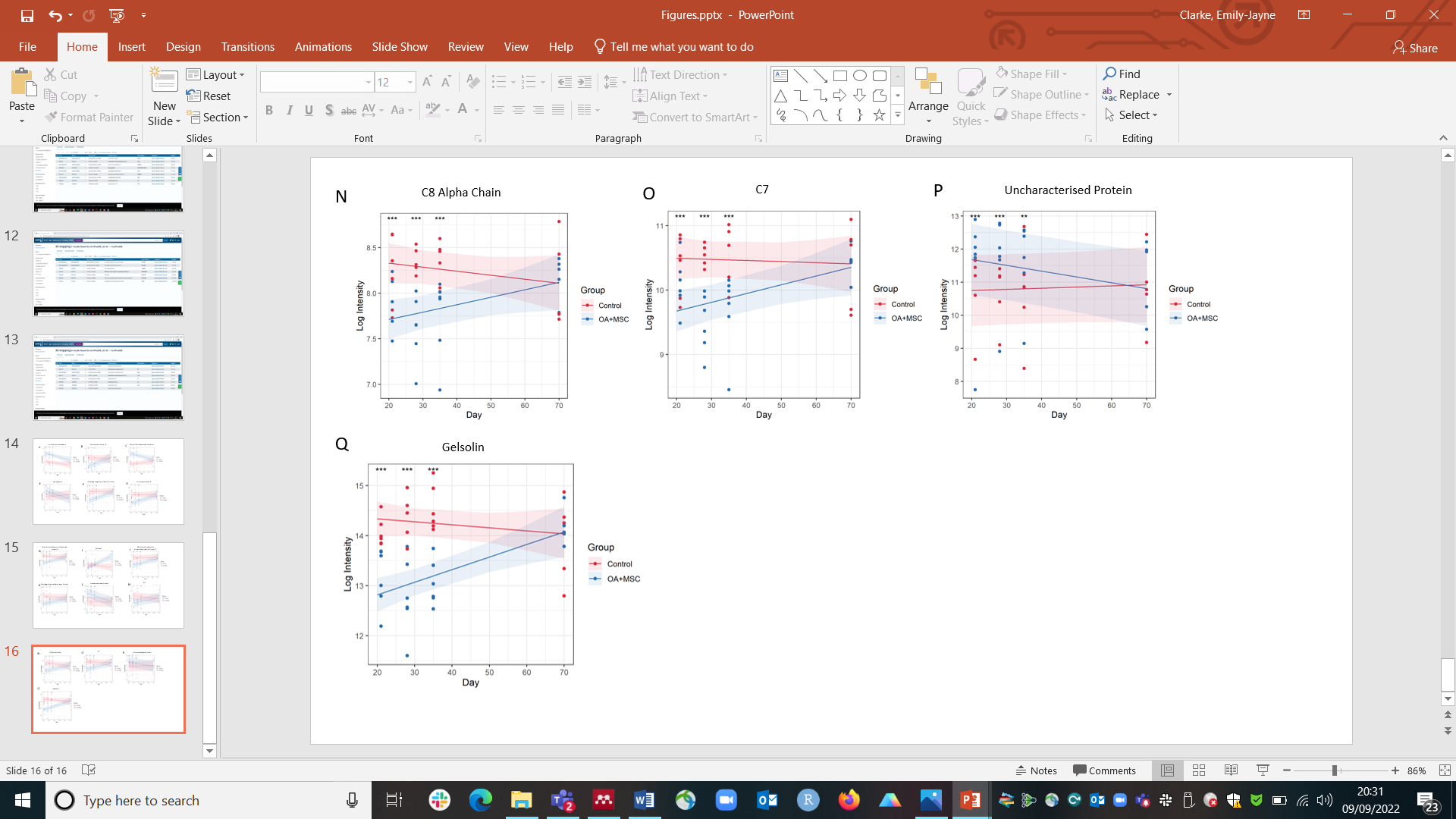
between experimental group (treatment OA+MSCs (treatment) and control over time (day 21, 28, 35 and 70), visualizing the expression change longitudinally. Pairwise comparisons were conducted post linear mixed model application to compare control and OA +MSCs protein expression per time point. Significance level, as determined by the generated FDR corrected P Value is shown using (P<0.05, *; P<0.01 **: P<0.001, ***, P<0.0001, ****).
